# A unique GCN5 histone acetyltransferase complex controls erythrocyte invasion and virulence in the malaria parasite *Plasmodium falciparum*

**DOI:** 10.1101/2021.02.03.429532

**Authors:** Jun Miao, Chengqi Wang, Amuza Lucky, Xiaoying Liang, Hui Min, Swamy Rakesh Adapa, Rays Jiang, Kami Kim, Liwang Cui

## Abstract

The histone acetyltransferase GCN5-associated SAGA complex is evolutionarily conserved from yeast to human and functions as a general transcription co-activator in global gene regulation. In this study, we identified a divergent GCN5 complex in *Plasmodium falciparum*, which contains two plant homeodomain (PHD) proteins (PfPHD1 and PfPHD2) and a plant apetela2 (AP2)-domain transcription factor (PfAP2-LT). To dissect the functions of the PfGCN5 complex, we generated parasites with the bromodomain deletion in PfGCN5 and the PHD domain deletion in PfPHD1. The two deletion mutants closely phenocopied each other, exhibiting significantly reduced merozoite invasion of erythrocytes and elevated sexual conversion. These domain deletions caused dramatic decreases not only in histone H3K9 acetylation but also in H3K4 trimethylation, indicating synergistic crosstalk between the two euchromatin marks. Domain deletion in either PfGCN5 or PfPHD1 profoundly disturbed the global transcription pattern, causing altered expression of more than 60% of the genes. At the schizont stage, these domain deletions were linked to specific downregulation of merozoite genes involved in erythrocyte invasion, many of which harbor the DNA-binding motifs for AP2-LT and/or AP2-I, suggesting targeted recruitment of the PfGCN5 complex to the invasion genes by these specific transcription factors. Conversely, at the ring stage, PfGCN5 or PfPHD1 domain deletions disrupted the mutually exclusive expression pattern of the entire *var* gene family, which encodes the virulent factor PfEMP1. Correlation analysis between the chromatin state and alteration of gene expression demonstrated that up- and down-regulated genes in these mutants are highly correlated with the silenct and active chromatin states in the wild-type parasite, respectively. Collectively, the PfGCN5 complex represents a novel HAT complex with a unique subunit composition including the AP2 transcription factor, which signifies a new paradigm for targeting the co-activator complex to regulate general and parasite-specific cellular processes in this low-branching parasitic protist.

**Author Summary:** Epigenetic regulation of gene expression plays essential roles in orchestrating the general and parasite-specific cellular pathways in the malaria parasite *Plasmodium falciparum*. Using tandem affinity purification and proteomic characterization, we identified a divergent transcription co-activator – the histone acetyltransferase GCN5-associated complex in *P. falciparum*, which contains nine core components, including two PHD domain proteins (PfPHD1 and PfPHD2) and a plant apetela2-domain transcription factor. To understand the functions of the PfGCN5 complex, we performed gene disruption in two subunits of this complex, PfGCN5 and PfPHD1. We found that the two deletion mutants displayed very similar growth phenotypes, including significantly reduced merozoite invasion rates and elevated sexual conversion. These two mutants were associated with dramatic decreases in histone H3K9 acetylation and H3K4 trimethylation, which led to global changes in chromatin states and gene expression. Genes significantly affected by the PfGCN5 and PfPHD1 gene disruption include those participating in parasite-specific pathways such as invasion, virulence, and sexual development. In conclusion, this study presents a new model of the PfGCN5 complex for targeting the co-activator complex to regulate general and parasite-specific cellular processes in this low-branching parasitic protist.

## Introduction

Packaging of the eukaryotic genomes with nucleosomes into chromatin affects all essential cellular processes such as transcription, DNA replication, and repair. A key mechanism for regulating chromatin structure involves post-translational modifications (PTMs) of nucleosomal histones, which can alter the accessibility of DNA and recruit distinct PTM readers and other effector proteins [1]. A multitude of histone PTMs such as acetylation, methylation, phosphorylation, ubiquitination, and sumoylation, act sequentially or combinatorially to create a “histone code” to facilitate or repress chromatin-mediated transcription [2–6]. Histone acetylation is a major PTM catalyzed by the histone acetyltransferase (HAT) enzymes and is correlated with active transcription [7]. HAT enzymes exist in large multimeric protein complexes such as the best-studied SAGA (Spt-Ada-Gcn5 acetyltransferase) complex [8] that is evolutionarily conserved from yeast to humans. The SAGA complex comprises 18–20 subunits, which are organized into functional modules including the HAT catalytic core, a histone deubiquitinase module, the TATA-binding protein (TBP) regulatory module, and the structural module [9]. In other SAGA-like complexes, the HAT catalytic core, consisting of the GCN5 acetyltransferase, ADA2, ADA3, and Sgf29, is conserved [9, 10]. Earlier studies suggested that SAGA regulates about 10% of genes in yeast and plants [11, 12], but a recent revisit of this issue in yeast revealed ubiquitous localization of SAGA at all gene promoters and reduced transcription of nearly all genes upon the disruption of SAGA [13, 14]. From these studies, SAGA appears to act as a general co-activator for all RNA polymerase II transcription, and its methyl reader (Sgf29) and acetyl reader (GCN5) subunits build synergistic crosstalk to coordinate transcription. As a co-activator complex that functions in the recruitment of the preinitiation complex, SAGA plays essential roles in metazoan development [15].

The human malaria parasite *Plasmodium falciparum* is an early-branching eukaryote, causing nearly half a million deaths in 2018 alone [16]. Its intricate lifecycle involving a vertebrate host and a mosquito vector requires precise regulation of transcription to cope with the comprehensive developmental program and environmental changes during host transitions [17–19]. Accumulating evidence indicates that the malaria parasite harbors unique properties of transcriptional regulation that are divergent from other eukaryotes even for the conserved transcription factors (TFs) [20–24]. Although the *Plasmodium* genome encodes the major components of the general transcription machinery, there is a general deficiency of specific *Plasmodium* TFs [25, 26]. Compared to the yeast *Saccharomyces cerevisiae* genome with ∼170 specific TFs [27], the similarly-sized genome of *P. falciparum* only has ∼30 TFs, including the 27 apicomplexan-specific AP2-domain TFs [26, 28, 29]. In contrast, the presence of almost a full complement of proteins involved in chromatin biology underlines the significance of epigenetic regulation in malaria parasites [23, 30]. One distinct feature of the *P. falciparum* epigenome is that it mainly consists of euchromatin with restricted heterochromatin regions at subtelomeres and a few internal loci [31–37]. The heterochromatin clusters are localized at the nuclear periphery and demarcated by high levels of H3K9me3 and binding of heterochromatin protein 1 (HP1), and they control antigenic variation, drug sensitivity, and gametocyte production [31, 32, 37–42]. In comparison, the *Plasmodium* euchromatin is characterized by low or no nucleosome occupancy at the transcription start sites (TSSs) and core promoters of highly expressed genes, which exhibits cyclic changes during the intraerythrocytic development cycle (IDC) [43–48]. Euchromatin is marked with the active histone PTMs such as H3K9ac, H4K8ac, and H3K4me3 [32, 49, 50], presumably deposited by the HAT enzymes PfGCN5 and PfMYST, and the methyltransferase PfSET1, respectively [51–54]. Of these euchromatic marks, H3K9ac at the promoter regions correlates well with the transcriptional status of the genes, whereas H3K4me3 shows stage-specific regulation and does not exhibit general correlation with transcription [50]. Despite the importance of the euchromatin structures as witnessed by the essence of the writers of these histone marks [53, 55], the mechanisms by which these active histone marks are deposited, maintained, and dynamically regulated during development are unknown. More intriguingly, since most of the genes encoding the diverse cellular pathways reside in the euchromatic regions, it is not clear how the cascade-like gene expression pattern is achieved.

The *P. falciparum* genome encodes a single GCN5 protein, PfGCN5, with a long, unique N-terminal extension lacking similarity to known protein domains, and a conserved C-terminal HAT enzyme domain that can acetylate histone H3 at K9 and K14 *in vitro* [51]. During the IDC, PfGCN5 is present as a full-length form, which also undergoes proteolytic processing by a cysteine protease-like enzyme [56]. PfGCN5 is essential for the IDC of the parasites; thus its function has been probed by chemical inhibition of its activity, which caused overall disturbance of transcription and gross reduction of H3K9ac, establishing a potential link between PfGCN5 and H3K9ac in the parasite [52, 57]. Recent efforts aiming to identify readers of the PTMs in *P. falciparum* led to the identification of putative PfGCN5-associated protein complex(es), which is highly divergent from the evolutionarily conserved SAGA complex [58]. Here, we used a tandem affinity purification (TAP) procedure to define a unique PfGCN5 complex and performed functional analyses of its key subunits. This work established the essential functions of this PfGCN5 complex in regulating cellular and metabolic pathways that are critical for parasite-specific processes such as antigenic variation, erythrocyte invasion, and sexual development.

## Results

### PfGCN5 Forms A Unique Complex Highly Divergent From The SAGA Complex

The evolutionarily conserved SAGA complex in eukaryotes comprises 18–20 subunits, which are organized into several modules including the HAT catalytic core consisting of GCN5, ADA2, ADA3, and Sgf29 [9, 10]. However, bioinformatic analysis of the *Plasmodium* genomes using conserved modular components of the SAGA complexes only identified two ubiquitous subunits GCN5 and ADA2 [51, 59], and a potential Tra1 homolog (PF3D7_1303800) [60], suggesting that the GCN5 complex(es) in these early-branching, parasitic protists might be highly divergent from the SAGA complex. Our recent work aiming to identify “readers” of modified histones with the H3K4me3 peptide surprisingly pulled down a putative PfGCN5 complex containing the PfGCN5, PfADA2, and two large proteins containing multiple plant homeodomains (PHDs) named PfPHD1 and PfPHD2 [58]. To precisely define the GCN5 complex(es) in *P. falciparum*, we tagged the C-terminus of the endogenous *PfGCN5* gene in the 3D7 strain with a PTP tag consisting of a protein C epitope, a tobacco etch virus (TEV) protease cleavage site, and two protein A domains (**Figure S1A, B**), which allows efficient TAP of protein complexes under native conditions with extremely low backgrounds [61, 62]. Integration-specific PCR confirmed the correct integration of the *PTP* tag at the *PfGCN5* locus (**Figure S1C)**. The transgenic parasites showed no noticeable *in vitro* growth defects (not shown). Western blot analysis using the anti-protein C antibody revealed that PfGCN5::PTP was expressed in all developmental stages of the IDC with the peak protein level in early trophozoites (**Figure S1D**). Six protein bands were detected and the band pattern agreed with that detected with an antibody against the PfGCN5 C-terminal fragment [56], confirming proteolytic processing of PfGCN5 (**Figure S1D**). Also, the processing of PfGCN5 was partially blocked with the cysteine proteinase inhibitor E64 (**Figure S1E**). Live-cell imaging of the green fluorescent protein (GFP)-tagged PfGCN5 parasite line [63] showed that the PfGCN5::GFP protein was expressed throughout the IDC and localized in the nucleus (**Figure S2**). Thus, we performed the TAP procedure using nuclear extracts from 10^9^ synchronized trophozoites of the PfGCN5::PTP parasite, which was followed by liquid chromatography and tandem mass spectrometry (LC-MS/MS) for accurate protein identification. The MS data were subjected to Significance Analysis of INTeractome (SAINT) using a threshold of probability above 94% and false discovery rate (FDR) below 1% [64].

Three independent experiments of TAP and LC-MS/MS consistently identified nine proteins (**Figure 1A**, **Table S1A, B**), presumably representing the core subunits of this PfGCN5 complex (**Figure 1A**). This is in sharp contrast to the detection of only some abundant cellular proteins in the control pulldown experiments (**Table S1A**). Seven proteins identified using the TAP procedure were also present in the PfGCN5-associated proteins identified by a single-step pulldown procedure [58]. In agreement with our earlier work showing interactions between PfGCN5 and PfADA2 [59], these two proteins were among the most enriched proteins in the PfGCN5::PTP pulldown, demonstrating the high efficiency of the TAP procedure and preserved integrity of the complex. Consistent with the recent PfGCN5::GFP pulldown [58], the PfGCN5 core complex includes two large proteins PfPHD1 (PF3D7_1008100) and PfPHD2 (PF3D7_1433400), each containing four PHD zinc fingers (**Figure 1A, B, S3A, B**). PfPHD1 also contains two AT hooks, which are DNA-binding domains with a preference for AT-rich regions [65]. Sequence analysis indicated that these PHDs belong to the PHD superfamily with some containing additional cysteine and histidine residues (called extended PHD, ePHD). The ePHD has been found to bind dsDNA, methylated H3K4, or other TFs [66–69]. Only the fourth PHD in PfPHD1 conforms to the canonical PHDs that bind to H3K4me3/2 [70] (**Figure S3C**). Our recent study confirmed that this domain indeed preferentially binds H3K4me3/2 [58]. Furthermore, these two proteins were found to harbor large numbers of acetylation sites in our acetylome study [71], indicating that they are substrates of protein lysine acetyltransferases. An AP2-domain family TF (PF3D7_0802100), named PfAP2-LT, which is highly expressed at the late stages of the IDC [29], was consistently identified in all experimental replicates of the pulldown studies. Of note, the interaction between the PfGCN5 N-terminal fragment and AP2-LT has been identified in a genome-wide yeast two-hybrid screen [72]. Further, a histone assembly protein PfNAPS (PF3D7_0919000) was also identified in the PfGCN5 interactome. Finally, in the PfGCN5 core complex are three hypothetical proteins with unknown functions (PF3D7_1019700, PF3D7_1364400, and PF3D7_1402800), which are conserved in all *Plasmodium* species (**Figure 1A, B**).

**Figure 1.**
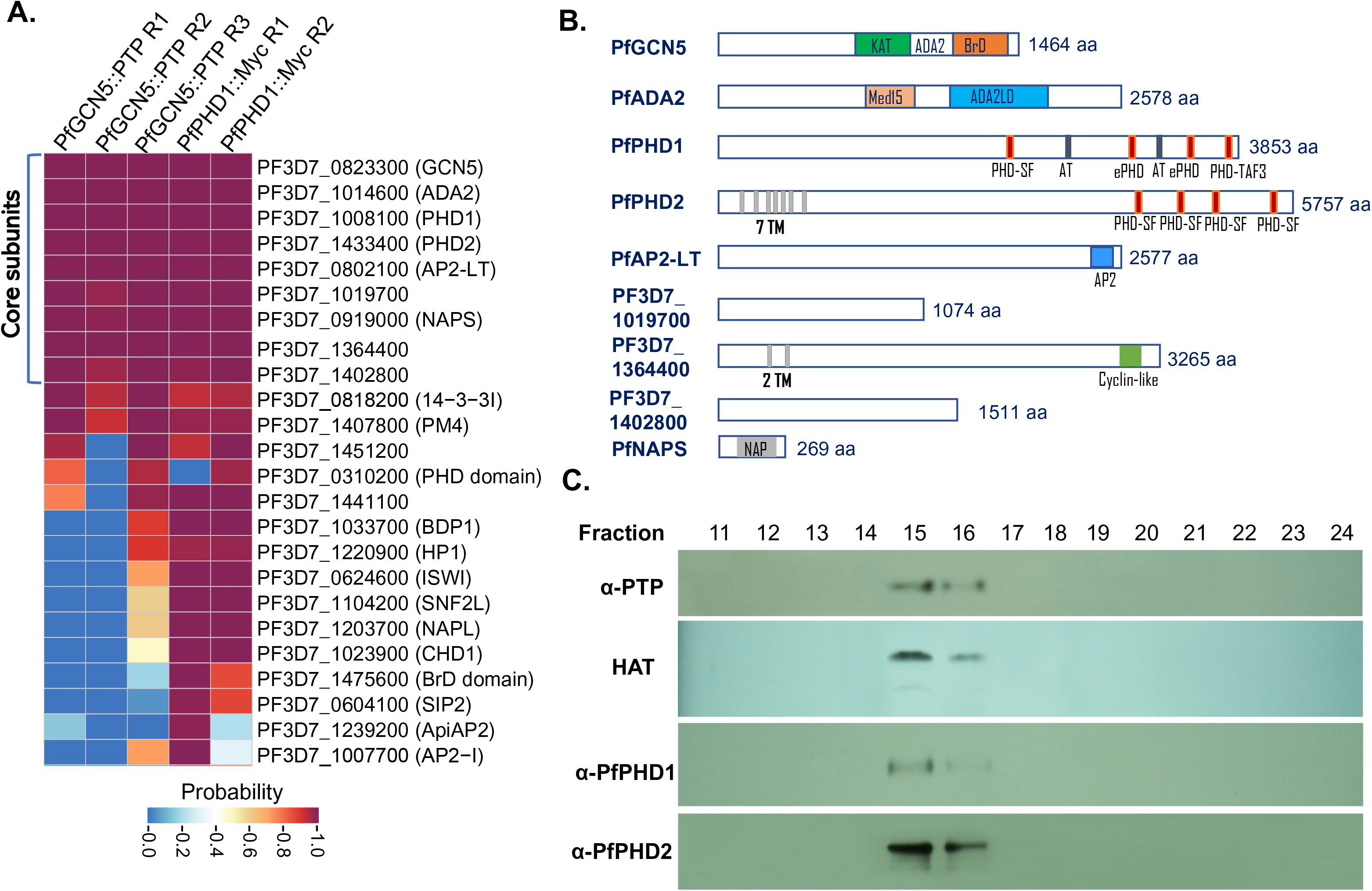
Identification of the PfGCN5 Core Complex in *P. falciparum*. (A) Proteins identified from parasite nuclear extracts by IP and by LC-MS/MS. TAP procedure was performed using the PfGCN5::PTP line (three replicates R1 – R3), while single-step IP with the anti-Myc beads was done using the PfPHD1::Myc parasite line (two replicates, R1, and R2). The proteomic data were analyzed by SAINT using a threshold of probability >94% and 1% FDR. Nine proteins consistently identified are marked as the PfGCN5 complex core subunits. Gene ID and annotation are shown on the right. (B) Schematic diagrams showing the features (putative domains and protein size) of the core subunits. (C) Gel filtration analysis of the PfGCN5 complex. Aliquots of different fractions were used for Western blots with anti-PTP, PfPHD1 and PfPHD2 antibodies, and for the HAT assay using recombinant histone H3.

To confirm that PfPHD1 and PfPHD2 are the core constituents of the PfGCN5 complex, rabbit polyclonal antibodies were generated against peptides of PfPHD1 (aa 3685-3702) and PfPHD2 (aa 5738-5756). Indirect immunofluorescence assay (IFA) showed that the pre-immune sera did not react with parasitized red blood cells (RBCs) (data not shown), whereas the anti-PfPHD1 and -PfPHD2 antibodies detected fluorescent signals in the parasite nuclei (**Figure S4A**). Nuclear extracts of trophozoites from the PfGCN5::PTP line were subjected to immunoprecipitation (IP) with anti-PfPHD1 and anti-PfPHD2 antibodies, and the immunoprecipitated proteins were analyzed by immunoblotting with the anti-protein C antibodies. In contrast to the control IP with the pre-immune sera, PfGCN5-PTP was only detected in the IP with the anti-PfPHD1 or -PfPHD2 antibodies, confirming co-purification of PfGCN5 with PfPHD1 and PfPHD2 (**Figure S4B**).

IP and proteomic analysis with the tagged PfPHD1 and PfPHD2 subunits suggests they may be present in different versions of the PfGCN5 complex [58]. Whereas IP from PfPHD1::3×HA only purified PfPHD1, PfGCN5, ADA2, and PF3D7_1402800, IP from PfPHD2::GFP identified 12 putative subunits with marginal enrichment of PfPHD1 but no pulldown of PF3D7_1402800 [58]. To clarify this discrepancy with our TAP results, we separately tagged two subunits in these two putative versions of the PfGCN5 complexes for reciprocal IP: the PfPHD1 with a C-terminal c-Myc tag and PF3D7_1019700 with a C-terminal GFP tag (**Figure S5**). Correct integration of the *c-Myc* tag at the *PfPHD1* locus and the *GFP* tag at the *PF3D7_1019700* locus was confirmed by Southern blot analysis and integration-specific PCR, respectively (**Figure S5A, B, S6A, B**). Nuclear localization of the PfPHD1::Myc and PF3D7_1019700::GFP proteins was verified by cellular fractionation–Western blot (**Figure S5C**) and live-cell imaging analysis (**Figure S6C**), respectively. Affinity purification of the trophozoite nuclear extracts from the PfPHD1::Myc parasites with the Myc-trap beads followed by LC-MS/MS consistently identified all 9 subunits of the putative PfGCN5 complex, including PfPHD2 (**Figure 1A, Table S1C, D**). Similarly, two IP replicates from the PF3D7_1019700::GFP identified 7 of the 9 core components of the PfGCN5 complex, including both PfPHD1 and PfPHD2 (**Table S1E, F**). These results are consistent with the presence of both PfPHD1 and PfPHD2 in the same PfGCN5 complex.

To estimate the size of the PfGCN5 complex, the purified native complex by TAP from the PfGCN5::PTP parasites was subjected to gel filtration, and fractions were assayed for HAT activity and Western blots using anti-PTP, -PfPHD1, and -PfPHD2 antibodies (**Figure 1C**). The results showed that PfGCN5 and its associated HAT activity, PfPHD1, and PfPHD2 all were detected in fractions 15 and 16, which is compatible with one major PfGCN5 complex in *P. falciparum* asexual stages. Based on the calibration using molecular mass standards, the size of the complex was approximately 2.3 MDa, which is comparable to the size (2.26 MDa) estimated based on the predicted molecular masses of the 9 core subunits (**Figure 1B**).

### Domain Deletions in PfGCN5 and PfPHD1 Cause Severe Growth Defects in Parasites

To characterize the function of the PfGCN5 complex in transcription regulation, we wanted to knock out the *PfGCN5, PfADA2, PfPHD1,* and *PfPHD2* genes by double-crossover homologous recombination but without success after multiple attempts (data not shown), indicating these genes are essential for parasite survival. This result is consistent with the mutagenesis scores in the genome-wide *piggyback* transposon mutagenesis study showing the essentiality of these genes [73]. Since PTM-binding domains are important for anchoring and withholding the respective proteins or complexes to the chromatin, we speculated that deleting these domains might disturb histone modifications without causing lethality to the parasite. Thus, we attempted to delete the bromodomain (BrD) and the PHD from the C-termini of PfGCN5 and PfPHD1, respectively, using a single-crossover gene disruption strategy, and meanwhile tag the C-termini of these truncated proteins with a GFP tag for sorting parasites with truncated PfGCN5 or PfPHD1 (**Figure S7A, C**). After transfection, the parasites were selected with WR99210, and GFP-positive parasites were cloned by sorting GFP-positive parasites using flow cytometry. Correct integration of the plasmids at the *PfGCN5* and the *PfPHD1* loci in the parasite genome was confirmed by Southern blots (**Figure S7B, D**). Phenotypic analyses of the parasites with the domain deletions in these two proteins revealed that the parasites with PfGCN5 BrD deletion (GCN5-ΔBrD) and parasites with PfPHD1 PHD domain deletion (PHD1-ΔPHD), to the greatest extent, pheno-copied each other (**Figure 2**). Both the GCN5-ΔBrD and PHD1-ΔPHD parasites grew significantly more slowly than the wild-type (WT) parasites; they only reached ∼1% parasitemia on day 7 as compared to ∼10% in WT parasites (**Figure 2A**). A more detailed analysis of the growth defects in these domain deletion lines showed that mature schizonts in these mutant parasites produced similar numbers of merozoites as the WT parasites (**Figure 2B**), but these merozoites had substantially reduced efficiency (by almost 80%) in the invasion of RBCs (*P <* 0.05, paired Wilcoxon test; **Figure 2C**). In addition, these domain deletion mutants also had a 2–3 h longer IDC than the WT parasites (*P <* 0.05, paired Wilcoxon test; **Figure 2D**), and a more extended ring stage (**Figure 2E, F, S6E**). Further, these domain deletion parasites were inclined to produce more gametocytes than WT (*P <* 0.05, paired Wilcoxon test; **Figure 2G**).

**Figure 2.**
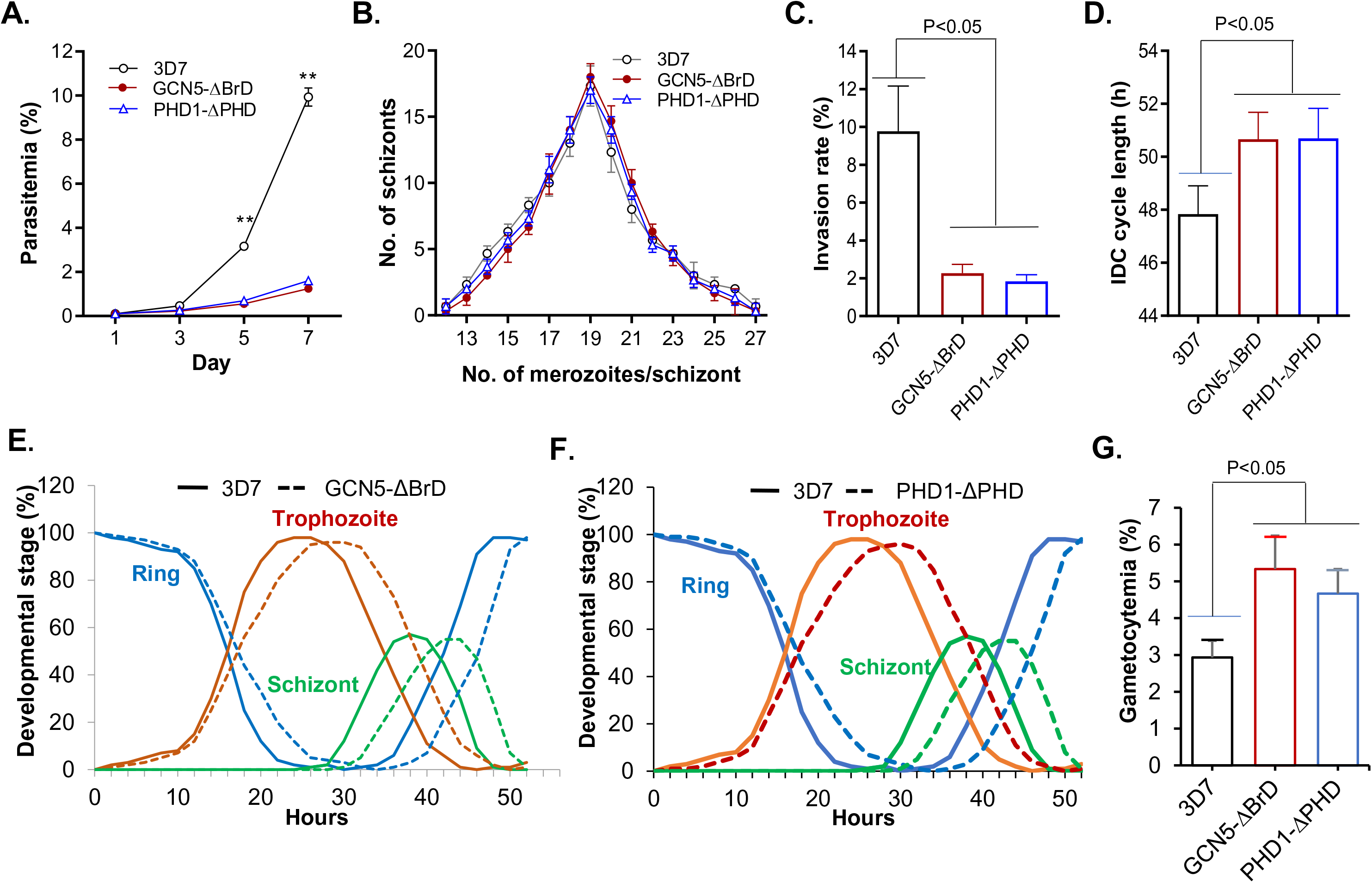
Growth phenotypes in PfGCN5 and PfPHD1 domain deletion mutants. (A) Asexual growth rates of WT 3D7, PfGCN5-ΔBrD::GFP, and PfPHD1-ΔPHD::GFP. ** indicate *P* < 0.01 (ANOVA) at days 5 and 7. (B) The distribution of the number of mature schizonts with a variable number of merozoite. No differences were identified among the three parasite lines (*P* > 0.05, ANOVA). (C) Merozoite invasion rates showing significantly reductions in GCN5-ΔBrD::GFP and PHD1-ΔPHD::GFP parasite lines (*P* < 0.05, paired Wilcoxon test). (D) The duration of the IDC showing significantly increased lengths in GCN5-ΔBrD::GFP and PHD1-ΔPHD::GFP parasite lines (*P* < 0.05, paired Wilcoxon test). (E, F) Detailed analysis of the IDC showing extended ring stage in the GCN5-ΔBrD::GFP (E) and PHD1-ΔPHD::GFP (F). (G) Gametocytemias at day 6 after induction of gametocytogenesis showing significantly increased gametocytemia in the two domain deletion mutants (*P* < 0.05, paired Wilcoxon test).

### Domain Deletions in PfGCN5 and PfPHD1 Are Associated with Globally Reduced H3K9ac and H3K4me3

*P. falciparum* has an extensively euchromatic epigenome with a preponderance of the histone marks H3K9ac and H3K4me3 [32, 50]. The presence of BrD and PHD in the PfGCN5 complex, which bind to acetylated H3K9/14 and H3K4me3/2 marks, respectively, strongly suggests that both domains may be required for anchoring the PfGCN5 complex to chromosomal regions to reinforce the euchromatic state. To determine the impacts of BrD and PHD deletions on the overall euchromatic histone marks, histones were purified from parasites of different developmental stages, and several histone marks were analyzed by Western blots. Consistent with PfGCN5 being the major HAT mediating H3K9 and H3K14 acetylation, deletion of BrD in PfGCN5 led to a significant reduction of the H3K9ac and H3K14ac levels, but the effect was more pronounced in the trophozoite stage, corresponding to the time of peak PfGCN5 expression (**Figure 3A**). Similarly, deletion of the PHD in PfPHD1 also resulted in the reduction of H3K9 and H3K14 acetylation. In comparison, domain deletions in PfGCN5 and PfPHD1 did not cause noticeable changes in H4 tetra-acetylation (at positions H4K5, 8, 14, and 20), which is mediated by another HAT protein, PfMYST [53]. Interestingly, domain deletions in these two subunits of the PfGCN5 complex also resulted in significantly reduced levels of H3K4me3 in trophozoites (**Figure 3A**), another major euchromatin mark conferred by the PfSET1 histone methyltransferase, highlighting the presence of extensive crosstalk between the two euchromatin marks. This result echoes well with findings from studies of the SAGA complexes in model organisms, where GCN5 deletion or the Sgf29 Tudor domain deletion reduced the levels of both H3K9ac and H3K4me3 [13, 14, 74]. Taken together, these results indicate that both BrD in PfGCN5 and PHD in PfPHD1 are important for anchoring the PfGCN5 complex to maintain the euchromatic histone marks.

**Figure 3.**
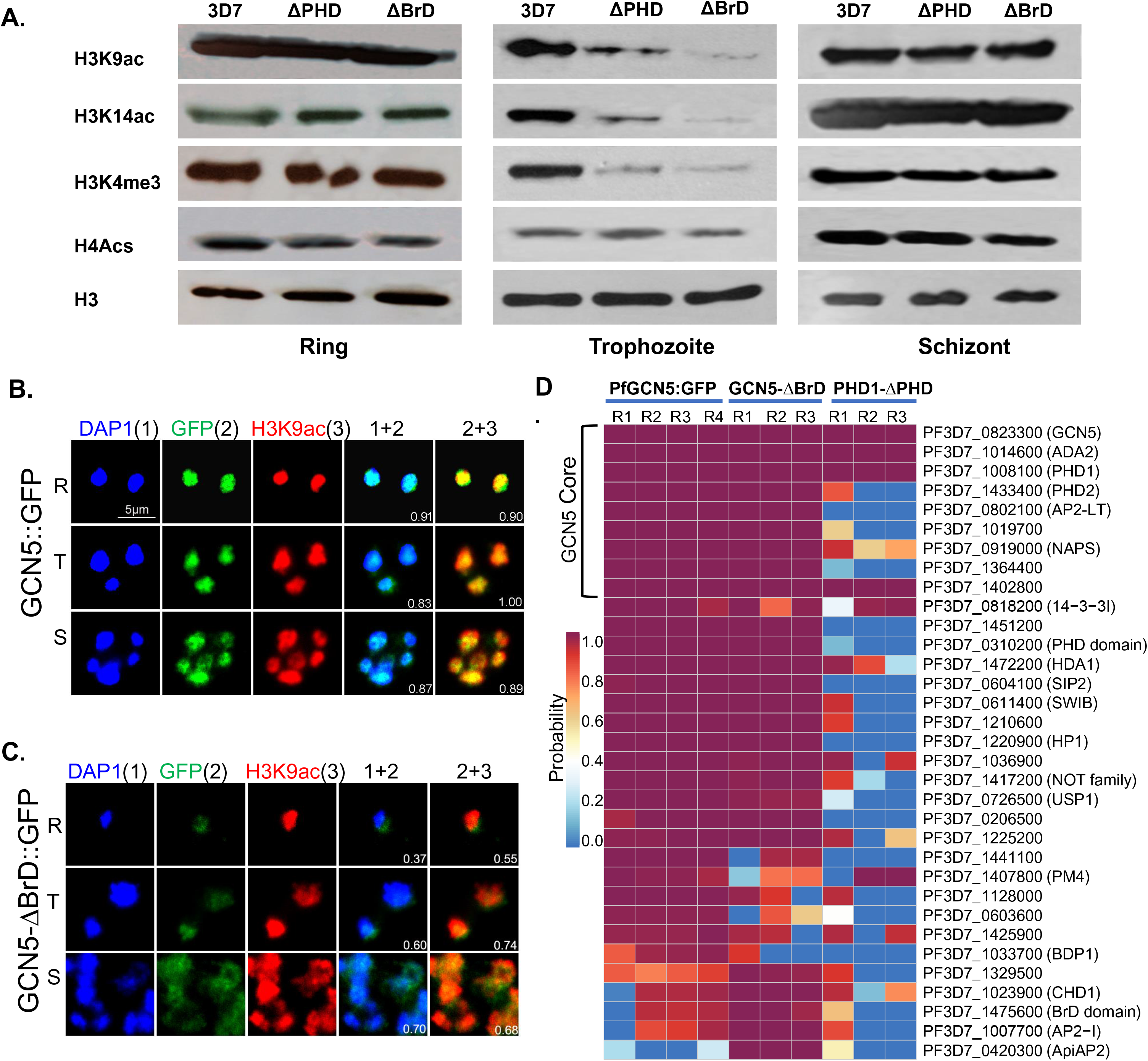
Domain deletions affect the abundance and localization of active histone marks and the integrity of the PfGCn5 complex. (A) The levels of active histone marks in 3D7, PfGCN5-ΔBrD::GFP (ΔBrD), and PfPHD1-ΔPHD::GFP (ΔPHD) parasite lines. Histones were purified from the ring, trophozoite, and schizont stages, and detected by Western blots with specific antibodies against the modified histones H3K9ac, H3K14ac, H3K4me3 and H4Acs. Anti-H3 antibodies were used for loading control. (B, C) Co-localization of full-length PfGCN5 (GCN5::GFP) (B) or truncated PfGCN5 (GCN5-ΔBrd::GFP) (C) with H3K9ac and DAPI by IFA with anti-GFP and H3K9Ac antibodies. Note the expansion of the truncated PfGCN5-ΔBrD::GFP and H3K9ac beyond the periphery of the euchromatin areas demarcated by DAPI staining. (D) Effects of domain deletions in PfGCN5 and PfPHD1 on complex integrity. Proteins were pulled down from the trophozoite nuclear extracts of the PfGCN5::GFP, GCN5-ΔBrD::GFP, and PHD1-ΔPHD::GFP parasite lines and identified by LC-MS/MS. R1, R2, R3 and R4 indicate individual repeats of the experiment. Shown here are proteins passing the threshold of SAINT (probability >94% and FDR <1%). The nine PfGCN5 complex core subunits were all detected in the IPs of PfGCN5::GFP and GCN5-ΔBrD::GFP, whereas only four of the core subunits were identified in the IPs of PHD1-ΔPHD::GFP.

Spatial compartmentalization of chromatins in the nucleus is critical for gene regulation in malaria parasites. The active chromatin marks H3K9ac and the H3K14ac overlap extensively with the DAPI staining (often used to define the euchromatin domains), whereas the heterochromatin mark H3K9me3 mostly occupies the nuclear periphery outside of DAPI [31, 75]. To analyze whether these changes in histone modifications were associated with altered spatial organization of the chromatins, we used IFA and live microscopy to observe the nuclear locations of these histone modifications as well as the truncated PfGCN5 and PfPHD1. In the PfGCN5-ΔBrD::GFP parasites, Western blot with the anti-GFP antibodies detected reduced expression of the truncated PfGCN5 protein, while the pattern of the PfGCN5 fragments remained similar (**Figure S8A**). Live microscopy of the GFP-tagged truncated PfGCN5-ΔBrD::GFP and PfPHD1-ΔPHD::GFP parasites showed that the GFP signals, while overlapping largely with the parasite nuclei from DAPI staining (**Figure S8B, C**), were more diffused than those in the WT parasites (**Figure S2**). Consistent results were obtained by IFA with anti-GFP and H3K9ac antibodies in WT and PfGCN5-ΔBrD::GFP parasites (**Figure 3B, C**). In the PfGCN5::GFP parasites, there were high levels of overlaps between the DAPI and PfGCN5 (*r*^2^=0.83–0.91), and between PfGCN5 and H3K9ac (*r*^2^=0.89–1.0), indicating that PfGCN5 is tightly associated with H3K9ac in the active euchromatin area demarcated by DAPI staining (**Figure 3B**). However, the levels of the co-localization substantially decreased in the PfGCN5-ΔBrD::GFP parasites (*r*^2^=0.37–0.74) (**Figure 3C**). Some signals of the truncated PfGCN5 were localized beyond the DAPI area, suggesting that PfGCN5 might have spread to the perinuclear heterochromatic area. Altogether, these results indicated that domain deletions in PfGCN5 and PfPHD1 altered the nuclear distribution of the truncated proteins and reduced the levels of euchromatin marks in the parasites.

### PHD Deletion in PfPHD1 Affects the Integrity of the PfGCN5 Core Complex

To determine whether BrD deletion in PfGCN5 and PHD deletion in PfPHD1 affected the integrity of the PfGCN5 complex, we performed IP by using the GFP-Trap antibodies with nuclear extracts from the individual domain deletion lines and analyzed the affinity-purified proteins by LC-MS/MS. Four replicates of IP using the PfGCN5::GFP parasites as the positive control consistently detected the core components of the PfGCN5 complex (**Figure 3D, Table S2A, B**). IP with the PfGCN5-ΔBrD::GFP parasites also confidently purified the 9 core subunits of the PfGCN5 complex, suggesting the deletion of the BrD from PfGCN5 did not affect the integrity of the complex (**Figure 3D, Table S2C, D**). However, only four major components of the PfGCN5 complex (PfGCN5, PfADA2, PfPHD1, and PF3D7_1402800) were detected from the PfPHD1-ΔPHD::GFP parasites (**Figure 3D, Table S2E, F**).

### BrD and PHD Deletions Profoundly Affect Global Transcription

Since GCN5-associated complexes facilitate transcription of target genes by bridging transcriptional activator and the preinitiation complex, deletion of domains from subunits of the complex that interact with histone tails weakens the anchoring and retention of the complex, leading to reduced transcriptional activation of the target genes [74, 76, 77]. To gain a mechanistic understanding of the PfGCN5 complex in transcriptional regulation, we compared the transcriptomes of the WT parasites and parasites with domain deletions. Parasites were highly synchronized from purified schizonts with a 3 h window, and RNA-seq analysis was performed during the IDC at 10, 20, 30 and 40 h after RBC invasion in the WT parasites and at 10, 23, 33 and 43 h in the parasites with BrD and PHD deletions to more closely match the developmental stages of the WT and domain-deletion parasites based on comparison of their IDC (**Figure 2E, F**). Compared to the phaseogram of WT parasites displaying a clear cascade-like gene expression pattern [18], PfGCN5 BrD deletion profoundly disturbed the global transcription pattern, causing 3533 (62.6%) genes to be differentially expressed in at least one of the four IDC time points analyzed (**Figure 4A**, **Table S3, S4**). Specifically, BrD deletion resulted in down-regulation of 997, 799, 861 and 902 genes, and up-regulation of 1127, 780, 846, and 368 genes at the ring, early trophozoite, late trophozoite, and schizont stage, respectively (**Figure 4B-D, S9A, B**). Noticeably, the numbers of up- and down-regulated transcripts were comparable at all stages except at the schizont stage, where 2.5-fold more transcripts were down-regulated than up-regulated (**Figure 4D**). In comparison, PHD deletion in PfPHD1 caused a similar but more profound disturbance of gene expression during the IDC, with 3870 (68.6%) transcripts being differentially expressed in at least one of the four stages analyzed (**Figure 4A**), which is congruent with the more substantial disruption of the PfGCN5 complex upon PHD deletion (**Figure 4**). The PfPHD1 PHD deletion resulted in the down-regulation of 872, 1021, 557, and 787 genes, and up-regulation of 1481, 1266, 648, and 1028 genes at the ring, early trophozoite, late trophozoite and schizont stage, respectively (**Figure 4D-F, S9C, D)**. Of note, only in late trophozoites did PfPHD1 PHD deletion disturb the expression of fewer genes than PfGCN5 Brd deletion (1205 vs. 1707 genes) (**Figure 4D**).

**Figure 4.**
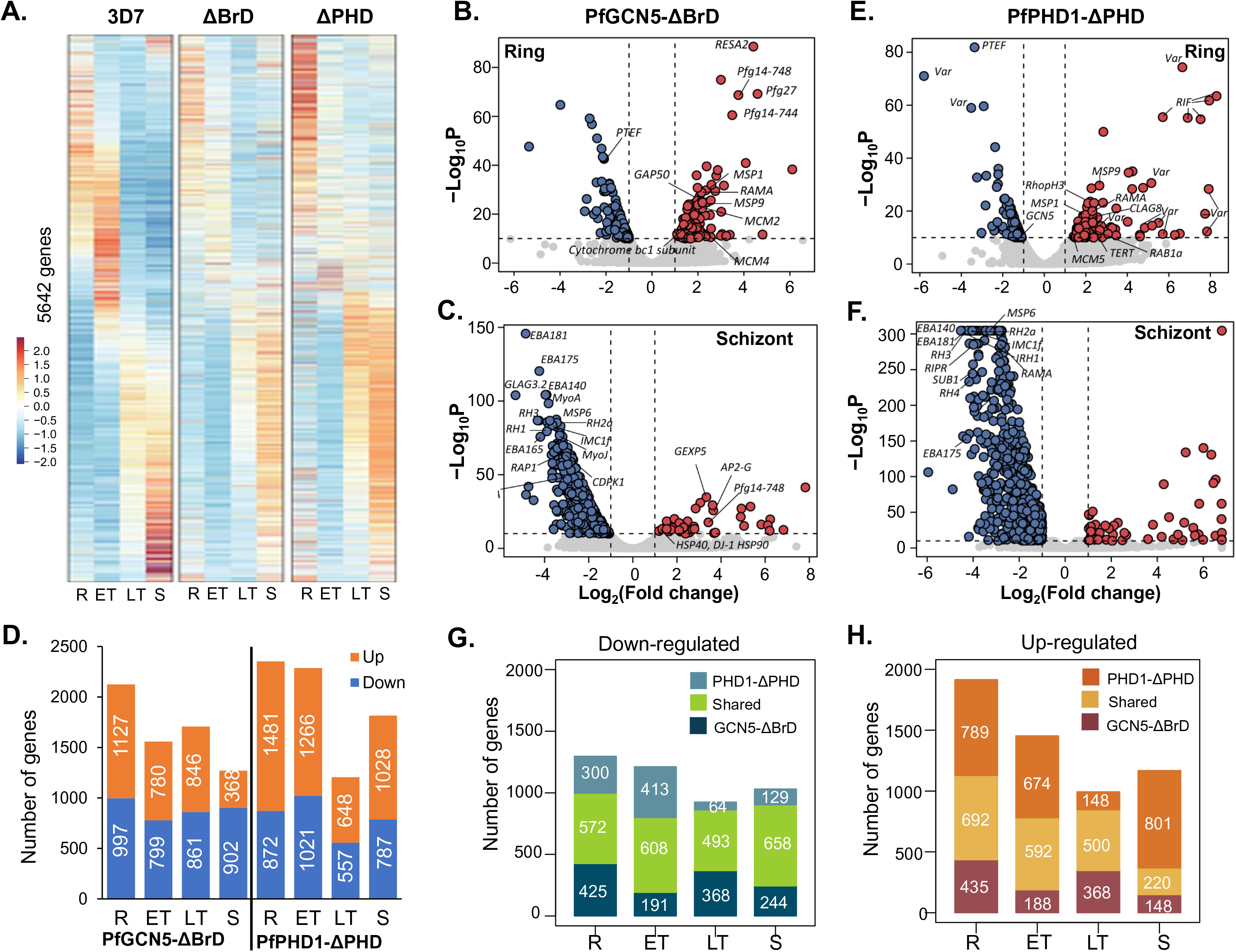
Global transcriptomic changes up domain deletions in PfGCN5 and PfPHD1. (A) The phaseograms of transcriptome from the WT 3D7, PfGCN5-ΔBrD::GFP (ΔBrd), PfPHD1-ΔPHD::GFP (ΔPHD) showing the disturbance of the cascade-like gene expression pattern in the deletion mutants at different developmental stages. R, ring; ET, early trophozoite; LT, late trophozoite; S, schizont. (B, C) Volcano plots showing altered gene expression at the ring (B) and schizont (C) stages in PfGCN5-ΔBrD::GFP compared to the WT 3D7. The x axis indicates log_2_ (Fold change) of the transcript level in PfGCN5-ΔBrD::GFP compared to WT 3D7, while the y axis indicates -log_10_ of the *P* values. (D) Number of genes with altered expression at different developmental stages of the IDC in the two domain deletion mutants. The up- and down-regulated genes are labeled in red and blue, respectively. (E, F) Volcano plots showing altered gene expression at the ring (E) and schizont (F) stages in PfPHD1-ΔPHD::GFP compared to the WT 3D7. The x axis indicates log_2_ (Fold change) of the transcript level in PfPHD1-ΔPHD::GFP compared to WT 3D7, while the y axis indicates log_10_ of the *P* values. (G, H) Overlaps of down-regulated (G) and up-regulated (H) genes between the PfGCN5-ΔBrD::GFP and PfPHD1-ΔPHD::GFP parasite lines at different stages.

With both PfGCN5 and PfPHD1 being integral members of the PfGCN5 complex, the global transcription changes resulted from the domain deletions of these two genes were remarkably similar; their transcriptomes showed a significant correlation between the respective stages with correlation coefficients ranging from 0.67 to 0.82 (**Figure S9E**). In addition, 44.1– 63.8% of the down-regulated genes in each stage were shared between the two domain deletion strains (**Figure 4G**). In comparison, 36.1–49.2% of the up-regulated genes were shared between the two domain deletion mutants in the ring, early and late trophozoites, but this up-regulated gene repertoire in the two domain deletion strains only shared 18.8% during the schizont stage (**Figure 4H**).

### BrD and PHD Deletions Affect Parasite-specific Cellular Pathways

To determine whether deletion of PfGCN5 BrD or PfPHD1 PHD affected specific biological processes, gene ontology (GO) enrichment analysis was performed on the genes with significantly altered expression (**Figure 5A, B, S10, Table S5**). During the early IDC stages, transcripts associated with cytoadherence, merozoite invasion, DNA replication, and organellar activities were up-regulated (**Figure 5A, S10A, Table S5**). Of particular interest, the 60 *var* genes, encoding the virulent factor PfEMP1 that mediates cytoadherence, were dramatically upregulated in the early stages, with their overall transcripts increased ∼4-fold upon BrD deletion and ∼25-fold upon PHD deletion (**Figure 5C, Table S6**). It is noteworthy that many *var* members were up-regulated, albeit *var2csa* was a major *var* gene expressed in PfGCN5-ΔBrD, suggesting activation of the overall *var* gene family. Western blot using antibodies against the conserved cytoplasmic ATS domain of the PfEMP1 proteins detected more complicated expression patterns and higher abundance of the PfEMP1 on the surface of RBC infected by trophozoite-stage parasites upon PfGNC5 BrD and PfPHD1 PHD deletions (**Figure 5D**). To further determine whether domain deletion in PfGCN5 and PfPHD1 activated multiple *var* members in a single infected RBC, we performed single-cell RNA-fluorescent in situ hybridization (FISH) using the type B *var* exon 2 as the probe [31], which is predicted to hybridize to 22 type B *var* genes. *Var* genes are clustered into 6-8 foci at the nuclear periphery and colocalized with the “telomere bouquets” [34], while the active *var* gene is localized to a *var*-specific expression site [78, 79]. Consistent with the mutually exclusive expression of *var* genes in single cells, the majority of the RNA-FISH positive cells contained one fluorescent spot (mean ± standard deviation, 1.04±0.21, n=67) indicating expression of a type B *var* (**Figure 5E**). In contrast, 43.9% and 58.2% positive rings had more than one hybridization signal in the PfGCN5-ΔBrD (1.55±0.66, n=66) and PfPHD1-ΔPHD parasites (1.91±0.98, n=79), respectively. In addition, 28.7% (45/157) of the WT 3D7 ring-stage parasites showed hybridization, which increased to 34.5% (61/177) and 41.3% (71/172) in the PfGCN5-ΔBrD and PfPHD1-ΔPHD rings, respectively. It is noteworthy that the hybridization signals were mostly localized in areas at the periphery of the DAPI staining. Thus, these results indicate the presence of multiple *var* expression sites in the PfGCN5-ΔBrD and PfPHD1-ΔPHD parasites.

**Figure 5.**
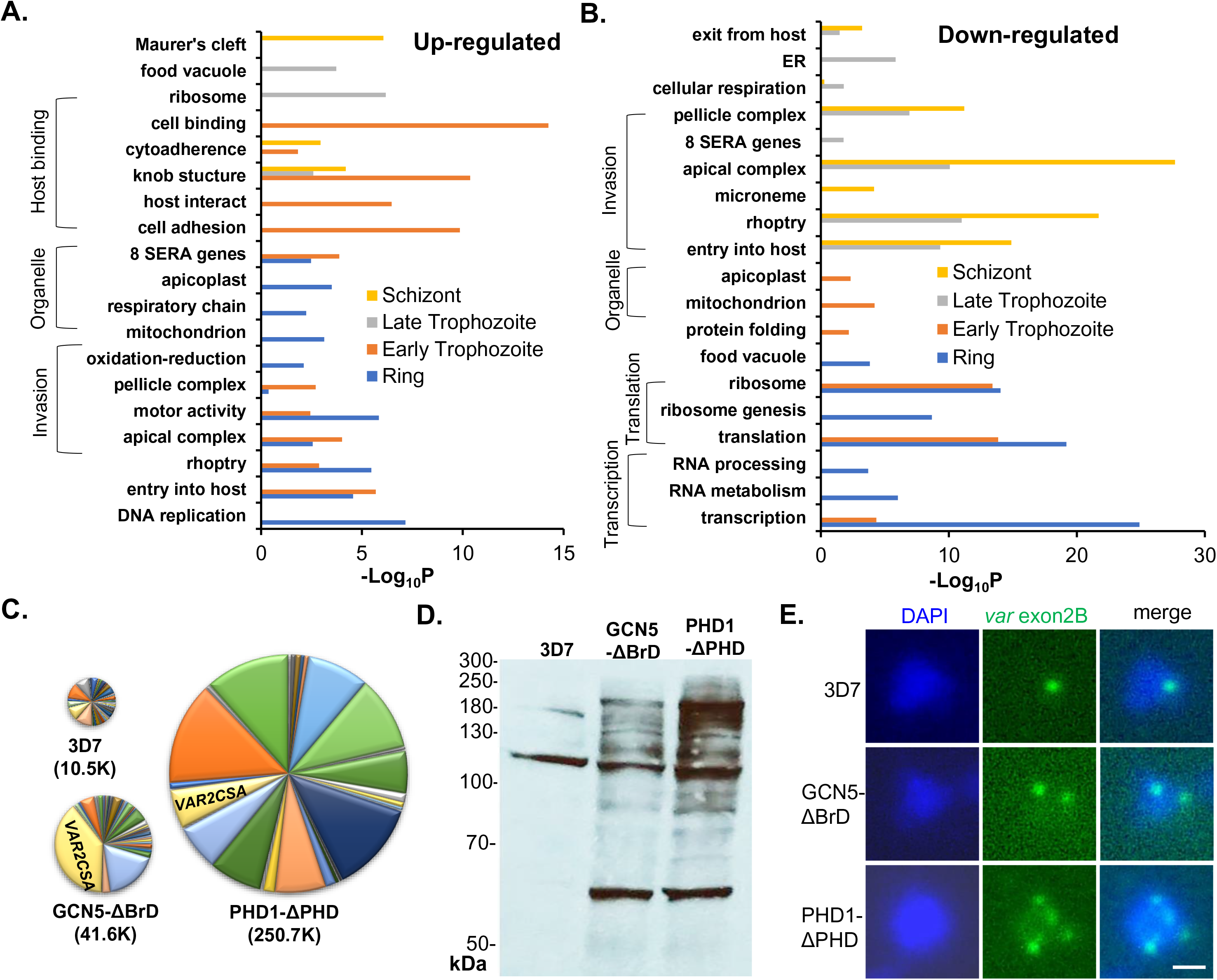
Biological processes and virulence gene expression altered upon domain deletions. (A, B) Gene ontology enrichment analysis of up-regulated (A) and down-regulated (B) genes in PfGCN5-ΔBrD::GFP parasites compared to WT 3D7. (C) Pie graphs showing the overall levels of the *var* gene transcripts in the WT 3D7, PfGCN5-ΔBrD::GFP, and PfPHD1-ΔPHD::GFP parasite lines at the ring stage. The numbers in parentheses are the total numbers of reads of all *var* genes identified by RNA-seq analysis. (D) Western blot showing PfEMP1 protein levels in the iRBC membranes of the WT 3D7, PfGCN5-ΔBrD::GFP, and PfPHD1-ΔPHD::GFP parasite lines with the anti-ATS antibodies. (E) Representative images of RNA FISH analysis showing single locus of the B-type *var* gene expression in the WT 3D7 and more than one B-type *var* gene locus in the two deletion mutants.

Conversely, genes in the biological processes of translation and transcription were significantly enriched in the down-regulated genes upon BrD or PHD deletion during the early IDC, which was probably responsible for the slowing down of development (**Figure 5B, S10B, C, Table S6**). During the late IDC (late trophozoites and schizonts), genes involved in RBC invasion were greatly reduced, which is consistent with the phenotype of reduced RBC invasion rates of the PfGCN5-ΔBrD and PfPHD1-ΔPHD merozoites (**Figure 5B, 6A, S10B, Table S6**). Of the 86 putative invasion-related genes [80], 76 showed peak expression at the late stages of IDC in WT parasites (**Figure 6A**). Except for *MSRP1* that was up-regulated, 75 genes were significantly down-regulated at the late stages in the PfGCN5-ΔBrD and PfPHD1-ΔPHD parasites (**Figure 6A**). These data collectively indicate the involvement of the PfGCN5 complex in the regulation of the general cellular processes such as transcription, translation, oxidoreduction, and organellar function, as well as parasite-specific processes of pathogenesis and host cell invasion.

**Figure 6.**
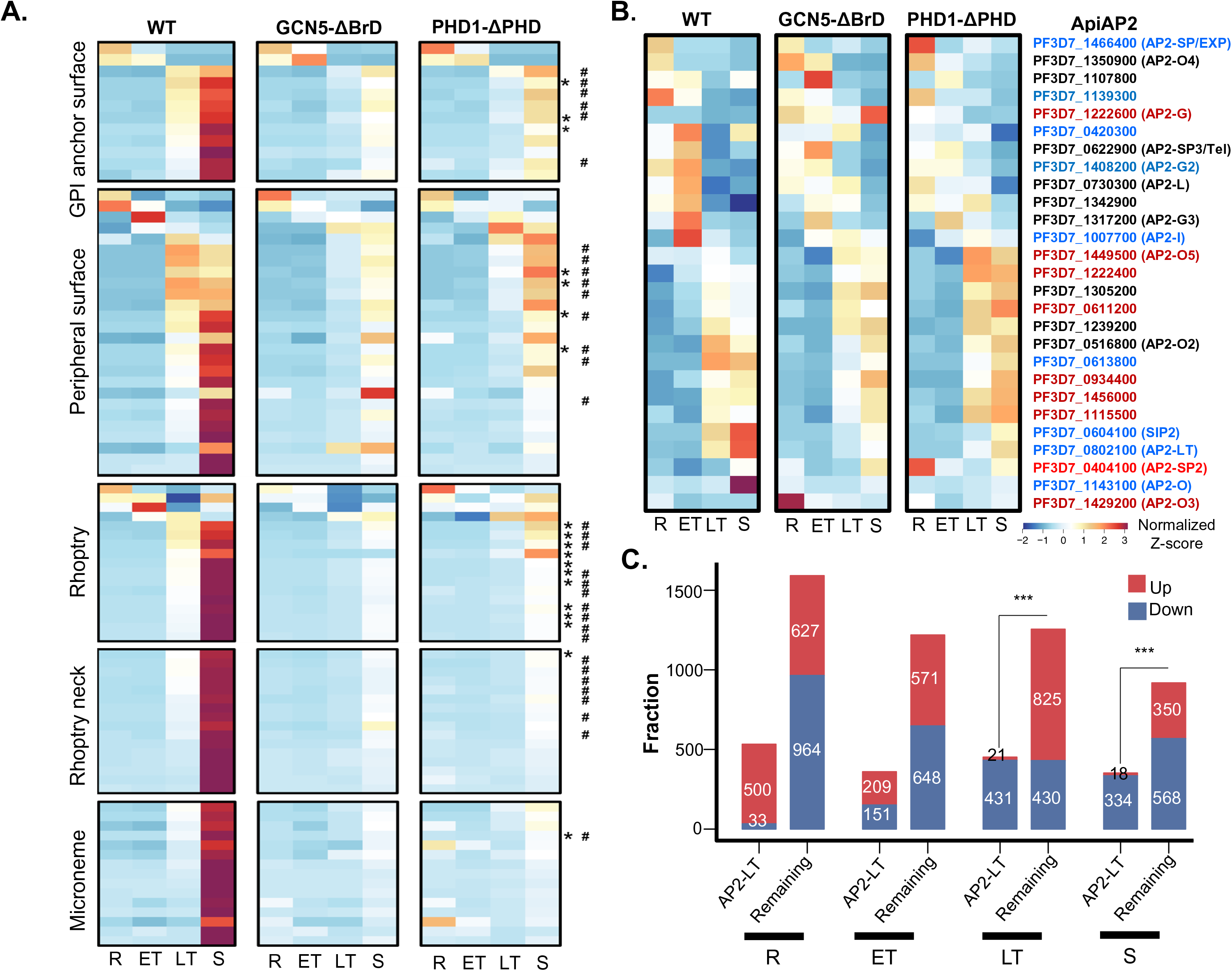
Down-regulation of invasion-related pathway and alteration of AP2 genes in domain deletion mutants. (A) Heatmaps displaying down-regulation of genes involved in the invasion of the RBC in the two deletion mutants. * and # indicate the AP2-I and putative AP2-LT target genes, respectively. (B) Heatmaps showing altered expression of the AP2 genes in the deletion mutants. (C) Putative AP2-LT target genes (with AP2-LT binding motifs) are significantly enriched in the down-regulated genes in PfGCN5-ΔBrD::GFP at the late stages. R, ring; ET, early trophozoite; LT, late trophozoite; S, schizont. ***, *P* <0.001 (Fisher’s exact test).

### Potential Coordination of the PfGCN5 Complex by AP2 Transcription Factors

The profound effects of PfGCN5 BrD/PfPHD1 PHD deletion on particular pathways such as invasion and cell adhesion suggest that these coordinated changes may involve the participation of specific TFs of the ApiAP2 domain family [81, 82]. The consistent pulldown with the PfGCN5 complex of the AP2-LT that is expressed abundantly in late stages of the IDC and occasionally other AP2 proteins (**Figure 1A**, **3D**) [58] suggests coordination of chromatin modification with transcription activators/repressors. To this end, we analyzed the transcriptional changes of all 27 ApiAP2 TFs after BrD or PHD deletion. Indeed, the cascade of AP2 transcriptions was substantially disturbed; some AP2-TFs such as AP2-SP2 and AP2-O3 were activated at stages when they are normally silenced in WT parasites, whereas some (e.g., AP2-O, AP2-I, and AP2-LT) were down-regulated at the stages when they are supposed to be active (**Figure 6B, Table S6**). PfSIP2 is associated with the chromosomal end clusters and is required for heterochromatin formation and genome integrity, including the silencing of subtelomeric *var* genes [83]. Its down-regulation upon BrD and PHD domain deletion may influence the organization of the subtelomeric heterochromatin, resulting in the overall de-repression of *var* genes (**Figure 6B**). The master regulator of gametocytogenesis AP2-G [84, 85] was consistently up-regulated upon Brd and PHD deletions (**Figure 6B**), which agrees with the increased gametocytogenesis detected in the mutant parasite lines (**Figure 2G**). The downregulation of AP2-I and AP2-LT might partially explain the downregulation of the invasion-related gene upon domain deletions. Of the 75 invasion-related genes down-regulated upon BrD or PHD deletion, 19 (mostly associated with the rhoptry) are targets of AP2-I, and 33 were predicted targets of AP2-LT (**Figure 6A, Table S6**) [29, 86]. The AP2-LT subunit of the PfGCN5 complex is predicted to bind to 986 genes with the motif sequence ACACA [29]. Analysis of the genes altered upon BrD deletion in PfGCN5 and PHD domain deletion in PfPHD1 revealed that genes down-regulated in late trophozoite and schizont stages were significantly enriched in those containing the AP2-LT binding motif (**Figure 6C, S10D**). This finding is consistent with the decreased recruitment of the PfGCN5 complex in the deletion mutants by AP2-LT to genes with AP2-LT binding motifs, resulting in extensive down-regulation of the gene categories during the late-stage development (**Figure 5B**). The unique presence of AP2 TFs in the GCN5 complexes in apicomplexan parasites suggests that the GCN5 complexes are specifically recruited to regulate the expression of certain clusters of genes [87].

### BrD and PHD Deletions Broadly Alter Chromatin Structures

The transcriptomic phaseograms showed that many genes were expressed out of the “phase” in the PfGNC5-ΔBrD and PfPHD1-ΔPHD parasites (**Figure 4A**); genes that are normally active were down-regulated, whereas genes supposed to be silent were active at the wrong time during the IDC. Since epigenetic regulation of gene expression in *P. falciparum* is most evident in heterochromatic regions [23], while gene expression from euchromatic regions correlates positively with the chromatin accessibility [48], we compared the chromatin status and accessibility of genes with altered expression upon PfGCN5 and PfPHD1 domain deletions. We first compared the up- and down-regulated genes with the accessibility of their promoters determined by the assay for transposase accessible chromatin sequencing (ATAC-seq) [48]. In all time points, the down-regulated genes upon BrD or PHD deletion are enriched in genes with more open chromatin structures at the promoters in the WT parasites, whereas up-regulated genes are significantly enriched in genes with less open promoters in the WT parasites (**Figure 7A**). Conversely, up-regulated genes upon BrD or PHD deletion are significantly more often associated with the heterochromatin loci that are normally enriched with HP1 and repressed during the IDC (**Figure 7B, Table S7**). This group of genes includes many variant gene families (*var*, *rifin* and *stevor*) and AP2-G (**Figure 5B, S10B, Table S7**). Among the genes that were up- regulated upon BrD or PHD deletion but have low accessibility in their promoters during the IDC in the WT parasites are the genes specific for sexual-stage development, which are normally silent during the IDC in the WT parasites. BrD deletion led to significant up-regulation of 353 gametocyte- and 401 ookinete-specific genes, respectively (**Figure 7C, D, Table S8**), and many were up-regulated at the ring stage. Similarly, PHD deletion caused up-regulation of 403 gametocyte- and 401 ookinete-specific genes, respectively (**Figure 7C, D, Table S8**). Among them, 151 gametocyte- and 199 ookinete-specific genes are shared between both deletional mutants. Taken together, both domain deletions similarly affected chromatin structures and led to the activation of genes involved in sexual development.

**Figure 7.**
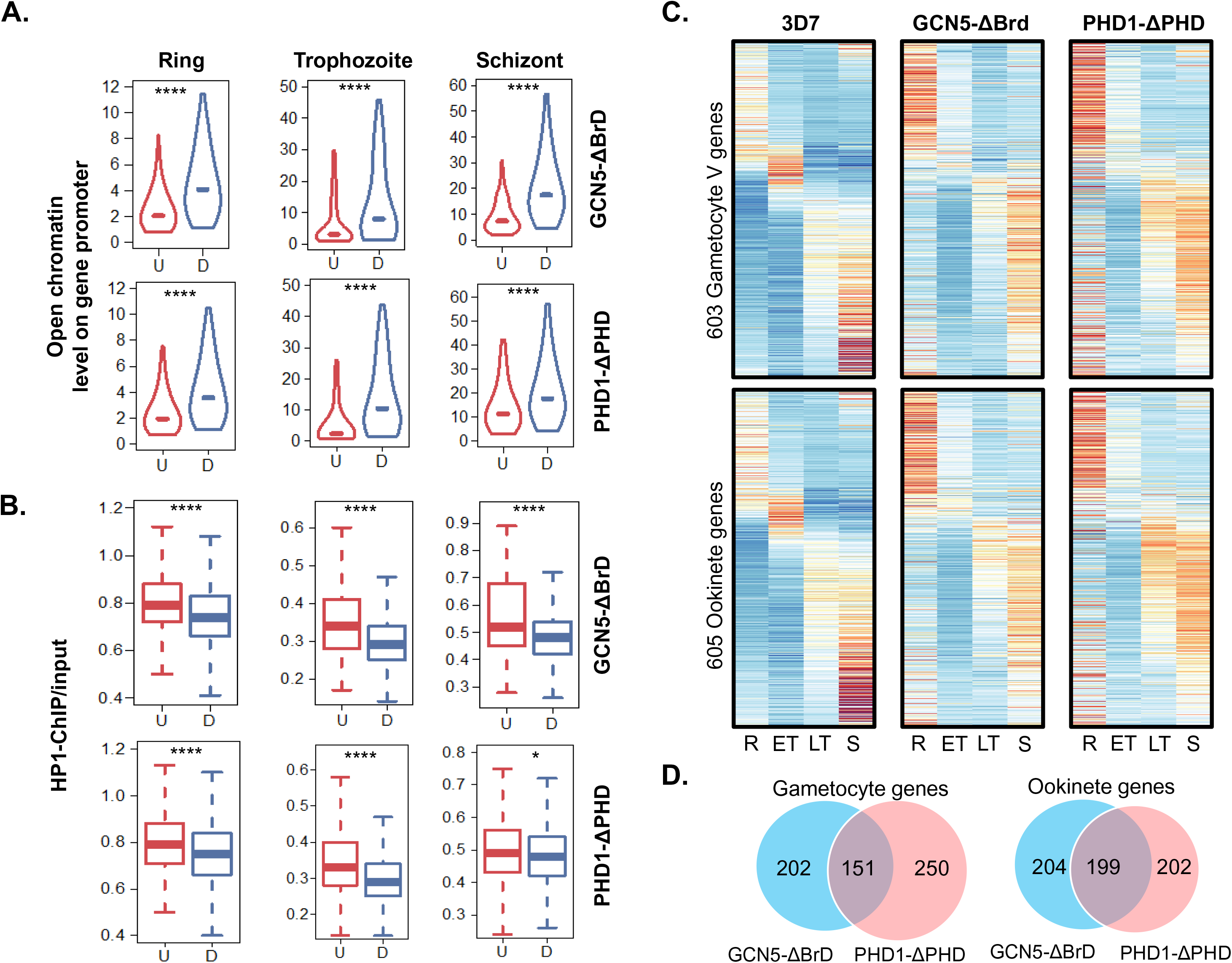
Correlation of genes showing altered expression in domain deletion mutants with promoter accessibility and chromatin states. (A, B) Changed levels of gene expression in PfGCN5-ΔBrd and PfPHD1-ΔPHD are negatively correlated with the accessibility of the promoters (from the ATAC-seq analysis) (A), but are positively correlated with the heterochromatin state (represented by the HP1 occupancy) (B). U, upregulation; D, downregulation. *, *P* <0.05; ****, *P* < 0.0001 (Wilcoxon rank-sum test). (C) Heatmaps displaying the transcriptional activation of gametocyte and ookinete genes in the domain deletion mutants. R, ring; ET, early trophozoite; LT, late trophozoite; S, schizont. (D) Overlaps of activated gametocyte- or ookinete-specific genes between PfGCN5-ΔBrd and PfPHD1-ΔPHD.

To verify that the PfGCN5 complex directly regulated the accessibility and chromatin state of the promoters, we selected three genes to evaluate the recruitment of the complex by chromosome immunoprecipitation (ChIP) followed by qPCR analysis. Compared to the WT parasites, the truncated PfGCN5-ΔBrD and PfPHD1-ΔPHD were significantly depleted at the *PfMSP1* promoter in schizonts, whereas they were enriched at the ring stage (**Figure S11A**). Their dynamic associations with the *PfMSP1* promoter were correlated with the down- and up-regulation of *PfMSP1* in these two domain deletion lines. Consistently, the PfGCN5-ΔBrD and PfPHD1-ΔPHD proteins were significantly enriched at the promoter of the *var2csa* gene and the gametocyte-specific gene *Pfg27/25* at the ring stage (**Figure S11B, C**), which was correlated with significant up-regulation of these two genes in the domain deletion parasites.

## Discussion

An unusual aspect of the chromatin-mediated gene regulation in the malaria parasite *P. falciparum* is that the parasite epigenome is dominantly euchromatic marked extensively with H3K9ac and H3K4me3 [32, 49], whereas heterochromatin-associated histone modifications H3K9me3 and H3K36me3 are localized to genes undergoing variable expression [31, 88]. This is contrasted to most eukaryotes where these heterochromatin marks are found in genes throughout the genome. Here we identified a unique GCN5 complex in *P. falciparum* responsible for depositing the euchromatic marks H3K9ac and H3K14ac, which is drastically different from the canonical SAGA complex that is conserved from yeast to human. Functional characterization of two major subunits demonstrated the crucial functions of the PfGCN5 complex in regulating global gene expression and parasite biology during its development in the host RBCs.

The evolutionarily conserved SAGA complex has a modular structure that supports multiple activities including histone acetylation, deubiquitination, and interactions with TFs [9]. Using multiple approaches, we refined the subunits of the GCN5 complex in *P. falciparum*, which completely lacks the deubiquitinase, the TBP-associated proteins, and the core structural module with the histone folds. Even for the HAT catalytic core [10], the *P. falciparum* GCN5 complex is also distinctive with the conservation of only the GCN5 and ADA2 homologs. Interestingly, the PfGCN5 complex also appears to differ substantially from the *Toxoplasma gondii* GCN5b complex [87, 89], as the latter contains additional proteins potentially involved in RNA binding and transcription elongation. In the PfGCN5 complex, the H3K4me3-binding activity, mediated by the tandem tudor domains of Sgf29 in the SAGA complexes [74, 90], is replaced by the PfPHD1 protein [58]. Moreover, PfPHD2 in the PfGCN5 complex contains four atypical PHDs, and may bind to other histone modifications. Importantly, since deletion of either BrD or PHD affected their localization and reduced the levels of both H3K9ac and H3K4me3, these domains are needed for anchoring and retention of the PfGCN5 complex on chromatin, similar to what was observed in the SAGA complex [13, 14, 74]. This result also implies the presence of synergistic crosstalk between the PfGCN5 complex and the histone H3K4me3 methyltransferase complex. As in model organisms, the binding of GCN5 BrD to H3K9Ac likely promotes H3 acetylation, which in return augments H3K4me3, since histone methyltransferases have a preference for the acetylated H3 tail [91–94]. The interactions between PfGCN5 and PfSET1 identified through yeast two-hybrid analysis further attests to the crosstalk between these euchromatic histone marks [72]. Given that the H3K4me3 levels are gradually increased toward the late stages of the IDC [50] and that most genes affected during PfSET1 knockdown are also expressed at the late stages [55], the intricate interplay between the writer complexes of these euchromatin marks needs to be further dissected.

Our earlier suggestion that PfPHD1 and PfPHD2 may represent different flavors of the PfGCN5 complex was based on the identification of only four proteins (PfGCN5, PfADA2, PfPHD1 and PF3D7_1402800) in the PfPHD1::3×HA pulldown and a complete lack of PF3D7_1402800 from the PfPHD2::GFP pulldown [58]. Here we provided evidence suggesting that both PfPHD1 and PfPHD2 are subunits of the same PfGCN5 complex. First, reciprocal pulldown with both the PfPHD1::Myc and PF3D7_1019700::GFP, which belong to the two putative PfGCN5 subcomplexes suggested earlier, identified the core components of the PfGCN5 complex with the abundant presence of both PfPHD1 and PfPHD2 (**Figure 1, Table S1**). Second, both PfPHD1 and PfPHD2 were co-eluted in the same fractions during gel filtration of the PfGCN5 complex. Third, the predicted size of the single PfGCN5 complex is compatible with the summation of the core subunits, while missing either of these two large PHD proteins would drastically reduce the size of the complex. Thus, this discrepancy may be due to the use of different tags for PfPHD1 (c-Myc vs 3×HA tag) and the different stringency of the analysis (1% FDR used here vs 10% used earlier). Also, we found that PfPHD1 could not be tagged with a larger tag such as the GFP (not shown), suggesting that tagging PfPHD1 with 3HA may interfere with the integrity of the PfGCN5 complex. Interestingly, pulldowns with both PfPHD1::3×HA and PfPHD1-ΔPHD::GFP identified the same four subunits of the PfGCN5 complex (**Figure 3D**). Thus, studies employing biochemical and cryogenic electron microscopy will allow a further resolution of the PfGCN5 complex.

The SAGA co-activator complex plays a critical role in regulating global gene expression [13, 14]. In *P. falciparum* with an unusual, dominantly euchromatic epigenome, the significance of the PfGCN5 complex is demonstrated by the essence of several core subunits for asexual development. Although domain deletion partially relieved the problem of lethality, deletion of either the BrD in PfGCN5 or PHD in PfPHD1 caused considerable growth defects during the IDC and altered expression of >60% of genes with an approximately equal number of up- and down-regulated genes. This defective gene expression pattern may be due to reduced levels of H3K9ac and H3K4me3, as well as mislocalization of the PfGCN5 complex. This may have led to the opposite changes in chromatin state, which are correlated with the genome-wide “out of phase” expression pattern. These global transcriptional changes are reminiscent of those when parasites were treated with the histone deacetylase inhibitor apicidin, which caused a reduction of H3K9Ac and H3K4me3 [95]. Analysis of genes with altered expression upon domain deletions for their chromatin state (HP1 occupancy) and promoter openness (ATAC-seq) provided indirect evidence supporting the mislocalization of the PfGCN5 complex in the deletion mutants. It is also noteworthy that PHD deletion in PfPHD1 caused more severe effects on gene expression, which can be explained by the disturbance of the integrity of the PfGCN5 complex upon PHD deletion, suggesting that PfPHD1 may play a scaffolding role for the structural integrity of the complex. Interestingly, the transcriptomic changes after BrD and PHD domain deletions (presumably due to reduced expression and mislocalization effects) are distinct from those after deletion of the SAGA subunits in yeast and human cells, which showed global downregulation of transcription [13, 14].

SAGA is recruited to the promoters through the interactions between Tra1 and TFs. Although a Tra1 homolog is present in the *P. falciparum* genome, it was not identified in the PfGCN5 complex. Another distinguishing feature of the PfGCN5 complex is the presence of an AP2-domain TF (PfAP2-LT) as a consistent member of the complex. Of note, the core GCN5b complex in the apicomplexan parasite *T. gondii* tachyzoites also contains multiple AP2 factors [87], suggesting a conserved characteristic in these lower-branching eukaryotes. These TFs would allow direct recruitment of the GCN5 complex to the target gene promoters, circumventing the need for a bridging factor such as Tra1. In addition to the identification of AP2-LT in the complex, two other AP2 factors (AP2-I and PF3D7_1239200) were also identified in the PfPHD1::Myc pulldown, suggesting that they might be either loosely associated with the GCN5 complex or represent additional flavors of the minor GCN5 complexes. In support of this notion, PfGCN5 was also identified in the AP2-I pulldown [86]. It is noteworthy that all pulldown experiments in this study were performed in the late trophozoite stage when AP2-LT is highly expressed, suggestive of the possibility that other AP2 factors may be associated with the PfGCN5 complex during different developmental stages. With the H3K9ac mark at the promoter regions dynamically following the pattern and level of transcription throughout the IDC [50], it is logical to propose that the dynamic recruitment of the PfGCN5 complex to different promoters is mediated by different AP2 factors. This hypothesis is compatible with the recruitment of the GCN5 complex by AP2-I to the promoters of invasion-related genes to acetylate the histones, which then recruit the BrD protein PfBDP1 [86]. In line with this, the genes predicted to be the targets of AP2-LT were mostly down-regulated upon PfGCN5 BrD and PfPHD1 PHD deletions, consistent with the AP2-LT-mediated recruitment of the PfGCN5 complex to the promoters of these genes. Of the 86 invasion-related genes expressed in merozoites, 33 were predicted to be the AP2-LT target genes, and 15 of these genes also are AP2-I targets (**Figure 6A**). Moreover, the majority of those AP2-LT target genes specifically affected by the domain deletions in PfGCN5 and PfPHD1 are expressed in late stages, coincidental with the peak expression of the AP2-LT (**Figure 6C**).

Antigenic variation in *P. falciparum* is mediated by the monoallelic expression of the ∼60 members of the *var* gene family [96]. *Var* gene clusters are located in the heterochromatin regions of the nuclear periphery and their expression is associated with the relocation and change of the promoter to a euchromatic state [38, 78, 97]. Such a mutually exclusive, monoallelic expression pattern of *var* genes is completely disrupted when the PfGCN5 BrD or PfPHD1 PHD domain was deleted, evidenced by the simultaneous expression of multiple *var* genes in single infected RBCs. This phenotype is similar to what was observed when the histone deacetylate PfHda2, PfSir2A, B, and the heterochromatin marker PfHP1 were experimentally knocked out or knocked down [98–101]. Given the presence of potential binding elements for AP2 factors in the *var* promoters, we cannot fully exclude the possibility that the activation of the *var* gene family upon PfGCN5 or PfPHD1 domain deletion may be due to altered recruitment of the PfGCN5 to *var* promoters through AP2 factors present in subsets of the GCN5 complex. However, the increased expression of most genes located in the heterochromatin regions marked by PfHP1 suggests that the expression of the whole *var* gene family may reflect a general loss of heterochromatin-based silencing instead of a specific effect on the mutually exclusive *var* gene expression. As the silent *var* genes cluster into 6-8 foci with the telomeres in the nuclear periphery, their activation upon PfGCN5 BrD or PfPHD1 PHD deletion did not seem to involve their “moving” to a single transcriptionally competent locus, but rather appearing as multiple active *var* loci in the nuclear periphery. This is consistent with the observed expansion of the truncated PfGCN5 or PfPHD1 to the outer nuclear compartments beyond the DAPI-stained central region. Moreover, the magnitude of all *var* transcripts compared with that in the WT parasite was significantly higher during PHD deletion (∼25-fold) than during BrD deletion (∼4-fold), which also agrees with the more severe effect of PHD deletion on the integrity of the PfGCN5 complex. This result emphasizes that maintaining the spatial organization of the different chromatin domains in *P. falciparum* is crucial for regulating antigenic variation.

This study has demonstrated the power of the TAP procedure for more precisely identifying protein complexes in malaria parasites. The nine subunits identified in this study may constitute the major PfGCN5 complex in the late stages of the IDC, while multiple variants of the PfGCN5 complex may exist to carry out different biological functions. Single-step IP identified large numbers of additional, less abundant proteins, which may represent those that are either associated less tightly with the PfGCN5 core complex or are the subunits of variant PfGCN5 complexes. In particular, a conserved histone modification reader protein Pf14-3-3I (PF3D7_0818200) was also identified in the pulldowns with both PfGCN5 and PfPHD1. Pf14-3-3I binds to purified parasite histones and H3 phosphopeptides [102] and its potential binding to H3Ser10p may further favor the recruitment of the PfGCN5 complex to acetylate H3K14 as identified in yeast [103], pointing to the presence of extensive crosstalk of epigenetic marks in *P. falciparum*. Other identified proteins with the GCN5 complex are mostly associated with the biology of chromatins including components of the SWI-SNF chromatin-remodeling complex and proteins with histone-interacting domains (e.g., BrD, WD40, PHD, CHD). Their potential associations with the PfGCN5 complex highlight the complexity of epigenetic regulation of gene expression in *P. falciparum*.

This study revealed a unique GCN5 complex in a lower eukaryotic parasite that is drastically distinct from the evolutionarily conserved SAGA complexes from yeast to human. The PfGCN5 complex, which is essential for regulating the stage-specific gene expression cascade, is also involved in the control of parasite-specific biologies such as RBC invasion and virulence. The critical role of the PfGCN5 complex in parasite biology and its significant divergence from the human host suggest that the PfGCN5 complex may be a vital target for chemotherapy against malaria parasites. Down this line, efforts are directed at identifying selective molecules inhibiting the GCN5 enzyme activity and selective inhibitors disrupting the interaction between the PfGCN5 BrD and acetylated histones [104, 105].

## Material and methods

### Parasite culture

The *P. falciparum* strain 3D7 and its genetically modified clones were cultured at 37°C in a gas mixture of 5% CO_2_, 3% O_2_ and 92% N_2_ with type O^+^ RBCs at 5% hematocrit in RPMI 1640 medium supplemented with 25 mM NaHCO_3_, 25 mM HEPES, 50 mg/L hypoxanthine, 0.5% Albumax II and 40 mg/ml gentamicin sulfate [106]. Synchronization of asexual stages was performed by two rounds of sorbitol treatment at the rings stage or by incubation of schizonts with RBCs for 3 h to obtain highly synchronized ring-stage parasites [107].

### Genetic manipulation of *PfGCN5* and its associated genes

To tag the C-terminus of PfGCN5 with the PTP tag [61, 62], the C-terminal *PfGCN5* fragment [nucleotides (nt) 3778-4758] was amplified from *P. falciparum* genomic DNA using primers F1 × R1. For the deletion of the PfGCN5 BrD and the PfPHD1 PHD domain, the *PfGCN5* fragment (nt 3286-4044) and the *PfPHD1* fragment (nt 3286-4044) were amplified using primers F2 × R2 and F3 × R3, respectively. All amplified fragments were first cloned into a modified pBluescript SK plasmid to fuse with the PTP or GFP and *pDT* 3’ UTR as described earlier [53, 108]. This cassette was then subcloned into pHD22Y at *Bam*HI and *Not*I sites to produce pHD22Y/PfGCN-PTP, pHD22Y/GCN5-ΔBrD-GFP, and pHD22Y/PHD1-ΔPHD-GFP, respectively. A similar strategy was used to tag PfPHD1 with c-Myc and PF3D7_1019700 with GFP. All primers used for tagging, domain deletion, PCR verification of integration and probe for Southern blot are listed in **Table S9A**.

Parasite transfection was done using the RBC loading method [109]. Briefly, 100 µg of plasmid were introduced into fresh RBCs by electroporation. Purified schizonts were used to infect the RBCs pre-loaded with the plasmid and selection was done with 2.5 nM of WR99210 for approximately 4 weeks with weekly replenishment of fresh RBCs until resistant parasites appeared. Resistant parasites were subjected to three cycles of drug on-off selection and single clones of parasites with stable integration of the constructs were obtained by limiting dilution [107]. For the parasites transfected with constructs containing a GFP tag, fluorescence-activated cell sorting was employed to clone the GFP-positive parasites. Correct integrations of plasmids into the parasite genome were screened by integration-specific PCR or Southern blot with the digoxigenin (DIG)-labeled probes using an established protocol [110].

### Purification of protein complexes

TAP was performed using the PTP-tagged PfGCN5 parasite line according to the published method [62]. Briefly, 10^9^ parasites were lysed in 5 volumes of the hypotonic buffer (10 mM HEPES, pH 7.9, 1.5 mM MgCl_2_, 10 mM KCl, 0.5 mM DTT, 0.5 mM EDTA) at 4°C for 10 min followed by centrifugation for 20 min at 500 × g. The resultant pellet (nucleus) was further lysed in 5 volumes of PA150 buffer (150 mM KCl, 20 mM Tris-HCl, pH 7.7, 3 mM MgCl_2_, 0.5 mM DTT, and 0.1% Tween 20) containing a protease inhibitor cocktail (Roche). The lysate was centrifuged for 10 min at 16,000 × g and the supernatant was incubated with 100 μl (settled volume) of IgG agarose beads (GE Healthcare) at 4°C for 2 h. The beads were washed twice with PA150 and equilibrated twice with the TEV buffer (150 mM KCl, 20 mM Tris-HCl, pH 7.7, 3 mM MgCl_2_, 0.5 mM EDTA, 1 mM DTT, and 0.1% Tween 20). To release the PfGCN5 and its associated proteins from IgG beads, the beads were incubated with 2 ml of TEV buffer containing 150 U of TEV protease and rotated overnight at 4°C. The supernatant was collected, and the beads were rinsed with another 4 ml of the PC150 buffer (150 mM KCl, 20 mM Tris-HCl, pH 7.7, 3 mM MgCl2, 1 mM CaCl_2_, 0.1% Tween 20). Then, 7.5 μl of 1 M CaCl_2_ were added to titrate the EDTA from the TEV buffer and the combined supernatant was incubated with the anti-protein C beads for 2 h at 4°C. The beads were washed four times with PC150 and eluted with the buffer containing 10 nM EGTA/5 mM EDTA. For the single-step pulldown of GFP-tagged or Myc-tagged protein, GFP- or Myc-trap (Cat# gta-20, RRID:AB_2631357 or Cat# yta-20, RRID:AB_2631369, Chromotek) beads were used with lysates from 10^9^ parasites according to the manufacture’s protocol.

### Mass spectrometry

The proteins in the elution were concentrated by Amicon Ultra centrifugal filters (Millipore Sigma) and separated briefly in a 10% Bis-Tris SDS-PAGE gel for 10 min. Proteins in gel were excised, in-gel digested, and analyzed by nano-LC/MS/MS using a Waters NanoAcquity HPLC system interfaced to a Q Exactive™ Hybrid Quadrupole-Orbitrap Mass Spectrometer (Thermo Scientific) [111]. Peptides were loaded on a trapping column and eluted over a 75 μm analytical column at 350 nL/min. MS and MS/MS were performed at 70,000 FWHM and 17,500 FWHM resolutions, respectively. The fifteen most abundant ions were selected for MS/MS. Parasite proteins were identified by searching the Uniprot *P. falciparum* protein database (v01/2014). Data were filtered at 1% protein and 0.2% peptide FDR, and at least two unique peptides per protein. Mascot DAT files were parsed into the Scaffold software for validation and filtering to create a non-redundant list per sample. The available mass spectrometry proteomics data have been deposited to the ProteomeXchange Consortium via the PRIDE [112] partner repository with the dataset identifier PXD023389 and 10.6019/PXD023389.

### Antibody generation

To generate antibodies against PfPHD1 and PfPHD2, a PfPHD1 peptide (DNGKLQKVDGRKKRRYHK, aa 3685-3702) and a PfPHD2 peptide (DDNVKAEDYKDENNDNDGD, aa 5738-5756) were synthesized and rabbits were immunized with these peptides. After three times immunizations, the antibodies were purified by affinity purification with peptides conjugated to the beads (Proteintech Group).

### Gel filtration

To access the size of the PfGCN5 complex, nuclear extract from the PfGCN5::PTP was incubated with IgG beads, eluted by TEV protease cleavage as described above, and applied to a Superose 6 gel filtration column (GE Healthcare). Molecular mass standards (Gel Filtration Calibration Kit HMW, GE Healthcare) were run under the same conditions to estimate the size of the complex. The fractions were analyzed by Western blotting using antibodies against the PTP tag, PfPHD1 and PfPHD2, while HAT activity in the fractions was measured as described previously [53].

### Growth phenotype analysis

The growth phenotypes of GCN5-ΔBrD::GFP and PHD1-ΔPHD::GFP lines during the IDC were compared with the WT 3D7 parasites as described [53]. To measure cell cycle progression, highly synchronous rings were obtained by incubation of schizonts with RBCs for 3 h. Progression of parasites through the IDC was monitored using Giemsa-stained smears every 2 h. Cycle time was determined as the duration between the peak ring parasitemias of two consecutive cycles. To measure parasite proliferation, synchronous cultures after two rounds of consecutive synchronization by sorbitol were initiated at 0.1% rings, and parasitemia was monitored daily for 7 days without replenishment of the RBCs. The number of merozoites produced per schizont was determined from mature segmenters. Three independent biological replications were done for each parasite line. Merozoite invasion assay was performed as described earlier [113]. The same numbers of purified merozoites from the WT, GCN5-ΔBrD::GFP and PHD1-ΔPHD::GFP lines were mixed with fresh RBCs, and the parasitemia of culture was determined 24 h later. The invasion rate was calculated as the percentage of merozoites invaded into RBCs. To measure the gametocyte development, gametocyte induction was conducted by using an established method [110, 114] and the gametocytemia was determined by counting gametocytes in Giemsa-stained thin blood smears at the middle developmental stage (stage III).

### Histone modifications

To estimate histone modifications in domain deletion mutants, histones were purified from the WT, GCN5-ΔBrD::GFP and PHD1-ΔPHD::GFP lines (Miao *et al.*, 2006). Equal amounts of the histones at each developmental stage were separated by 15% SDS/PAGE and transferred to nitrocellulose membranes. Western blotting was performed using a standard procedure with anti-acetyl histone H3, H3K9Ac (Catalog no. 07-352, RRID:AB_310544, Millipore), anti-tri methyl histone H3, H3K4me3 (catalog no. 07-473, RRID:AB_1977252, Millipore) and anti-acetyl histone H4, H4Ac (catalog no.06-598, RRID:AB_2295074, Millipore) at 1:1000 dilution as the primary antibodies and horseradish peroxidase-conjugated goat anti-rabbit IgG (diluted at 1:2000) as the secondary antibodies. The detected proteins were visualized using an enhanced chemiluminescence (ECL) kit (Invitrogen).

### Immunofluorescence assay (IFA)

IFA was performed as described [115, 116]. The parasitized RBCs were washed once with PBS and the cell pellet (∼100 μl) was fixed with 1 ml of 4% (v/v) paraformaldehyde and 0.0075% (v/v) glutaraldehyde in PBS for 30 min followed by 10 min quenching with 50 mM glycine in PBS. Fixed cells were washed twice with PBS and treated with 0.5% (v/v) Triton X-100 in PBS for 10 min. Then, cells were washed twice with PBS and blocked in 3% (v/v) BSA for 1 h at room temperature. The anti-PfPHD1 (1 μg/ml), PfPHD2 antibodies (1 μg/ml), goat anti-GFP (1:2000; ab6673; Abcam, RRID:AB_305643, USA) and rabbit anti-H3K9ac (1:1000; 06-942, RRID:AB_310308, Millipore, USA) antibodies in PBS containing 1% BSA were added and incubated for another 1.5 h. After washing the cells three times with PBS, FITC-conjugated goat anti-rabbit IgG antibodies (Cat# F6005, RRID:AB_259682, Sigma, USA), Alexa fluor® 488-conjugated secondary donkey anti-goat IgG antibody or Alexa fluor® 594-conjugated secondary goat anti-rabbit antibody IgG antibody (A32814 RRID:AB_2762838 and R37117 RRID:AB_2556545, Thermo Fisher Scientific, USA) were added at 1:2000 dilution in 3% (v/v) BSA and incubated for 45 min. Nuclear staining was performed by incubating slides with 4’,6-diamidino-2-phenylindole (DAPI, final 0.5 μg/mL; Invitrogen). Images were captured using an epifluorescence microscope (Nikon Eclipse Ni, USA; 100x/1.4 oil immersion lens) and were processed by Adobe Photoshop CS (Adobe Systems Inc. San José, CA). To quantitate co-localizations, images from at least 20 parasites were randomly selected, analyzed by ImageJ (1.52a; http://imagej.nih.gov/ij), and Pearson’s coefficients were calculated.

### Transcriptome analysis

To compare the transcriptomes during the IDC among the WT, GCN5-ΔBrD::GFP and PHD1-ΔPHD::GFP lines, RNA-seq was performed. Three replicates of total RNA from parasites at ring, early trophozoite, late trophozoite and schizont stages were harvested by using the ZYMO RNA purification kit, and used to generate the sequencing libraries using the KAPA Stranded mRNA Seq kit for the Illumina sequencing platform according to the manufacturer’s protocol (KAPA biosystems). Libraries were sequenced on an Illumina HiSeq 2500 in the Rapid Run mode using 100 nt single read sequencing. Reads from Illumina sequencing were mapped to the *P. falciparum* genome sequence (Genedb v3.1) using HISAT2 [117]. The coverage was analyzed by using the bedtools [118]. The expression levels and the differential expression were calculated by FeatureCounts and DESeq2 [119, 120] with the criteria of ≥ 2 fold of alteration and P-adjustment <0.01. The GO enrichment was performed on PlasmDB (https://plasmodb.org/plasmo/). RNA-Seq data were submitted to NCBI GEO repository (accession number GSE164070).

### Phaseogram of the transcriptomes of *P. falciparum* IDC

The sine wave model was utilized here to model the gene expression timing [121]. The gene transcription level from RNA-seq was first normalized as TPM (transcripts per million). Only the differential expressed genes in PfGCN5-ΔBrd or PfPHD1-ΔPHD as compared with WT were considered for the analysis. The TPM of each gene *E(t)* was modeled as

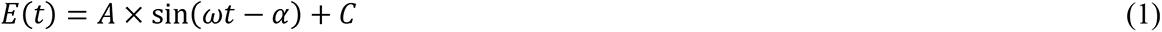

Where *E(t)* = [*TPM*_12h_, *TPM*_24h_, *TPM*_36h_, *TPM*_48h_] is the TPM at the *t =* [12, 24, 36, 48] hours of sample collection, *ω* is the angular frequency and given by *ω* = 2*π*/48, *A* is the amplitude of the expression profile, and *C* is the vertical offset of the profile from zero. To identify the parameter *α* and *A*, *A* × sin(*ωt* − *α*) are changed to

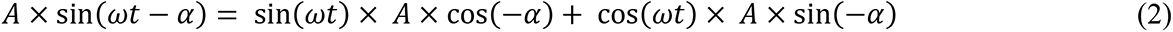

Then the R command *lm* was used to fit a linear regression model between *E*(*t*) and sin(*ωt*) + cos(*ωt*). The fitting coefficient from the *lm* result indicates *A* × cos(−*α*) and *A* × sin(−*α*). The *α* and *A* were calculated as

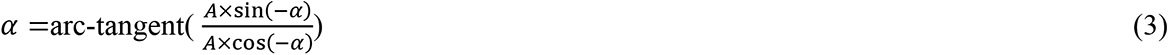

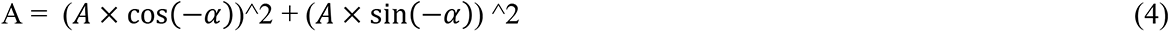

The *α* indicates the horizontal offset of the profile from zero, which is used in the phaseogram to order the gene in the heatmap.

### Association between chromatin structures and transcriptomic changes upon domain deletions

To quantify the association between open chromatin accessibility and transcriptional changes upon BrD deletion in PfGCN5 or PHD domain deletion in PfPHD1, we retrieved the ATAC-seq profile showing ATAC-seq peaks upstream the TSSs [122]. Each TSS was assigned to the nearest ATAC-seq peak with a distance restriction lower than 1 kb. The values of chromatin accessibility (ATAC-seq RPM + 0.1)/(gDNA RPM + 0.1) were then compared to the altered expression from the up- or down-regulated genes after domain deletions, where RPM represents the scaled reads per million reads. To investigate the association between the PfHP1 occupancy and transcriptional changes in the domain deletion mutants, the PfHP1 values (ChIP/input ratio) along the coding sequence were downloaded [123] and compared with the altered expression from the up and down-regulated gene after domain deletions.

### ChIP quantitative PCR

ChIP-qPCR was performed as described [52, 86, 124] with some modifications. Synchronized GCN5-ΔBrD::GFP and PHD1-ΔPHD::GFP parasite lines at the ring stage [10–16 h post-invasion (hpi), ∼5 × 10^9^ infected RBCs (iRBCs)] and schizont stage (40–46 hpi, ∼1.5 × 10^9^ iRBCs) were harvested and crosslinked with paraformaldehyde (1% final concentration; EMS, USA) at 37°C for 15 min with agitation and then immediately neutralized by adding glycine (0.125 M final concentration) on ice for 5 min with agitation. The fixed iRBCs were lysed with saponin (0.06% final concentration; Sigma, USA) on ice for 5–10 min. Parasites were treated with a lysis buffer (10 mM KCl, 0.1 mM EDTA, 0.1 mM EGTA, 1 mM DTT, 10 mM Hepes pH 7.9, 1 x Protease inhibitor) and then gently homogenized using a douncer to free nuclei. Pelleted nuclei were suspended in a shearing buffer (0.1% SDS, 5 mM EDTA, 50 mM Tris-HCl, pH 8.1, 1X Protease inhibitor) [124]. Sonication was performed using a rod bioruptor (Microson ultrasonic cell disruptor, Misonix, Inc. USA) at high power for 20 cycles of 30 sec ON/30 sec OFF, resulting in sheared chromatin of approximately 100–1000 bps. 50µl of input samples was set aside before the remaining chromatin was diluted in incubation buffer (0.01% SDS, 1.5% Triton X-100, 0.5 mM EDTA, 200 mM NaCl, 5 mM Tris-HCl, pH 8.1). Chromatin (75 µl/400 ng) was incubated with GFP-Trap® (Cat# gta-20, RRID:AB_2631357, ChromoTek, Germany) overnight at 4°C while rotating. Beads were then washed for 5 min at 4°C while rotating with the following: buffer 1 (0.1% SDS, 1% Triton X-100, 150 mM NaCl, 2 mM EDTA, 20 mM Tris HCl, pH 8.1); buffer 2 (0.1% SDS, 1% Triton X-100, 500 mM NaCl, 2 mM EDTA, 20 mM Tris HCl, pH 8.1), buffer 3 (250 mM LiCl, 1% NP-40, 1% Na-deoxycholate, 1 mM EDTA, 10 mM Tris HCl, pH 8.1) and finally twice with buffer 4 (10 mM EDTA, 10 mM Tris HCl, pH 8). The immunoprecipitated chromatin was eluted with the elution buffer (1% SDS, 0.1M NaHCO_3_) at room temperature for 15 min with rotation. The eluted chromatin and input samples were reverse cross-linked in 10% SDS, 1 M NaHCO_3,_ 5 M NaCl, 10% Triton X-100 at 45°C overnight while shaking and purified by the phenol:chloroform method. For qPCR, the concentration of immunoprecipitated gDNA was determined by Qubit dsDNA Broad-Range Assay Kit (Invitrogen, USA), and 10 ng per well in triplicate were used for qPCR using the FastStart™ Universal SYBR® Green Master [Rox] (Sigma-Aldrich, USA) as described [52]. Primer pairs targeting 5′UTRs were designed to amplify fragments less than 200 bp (**Table S9B**). Fold enrichment relative to constitutively expressed reference gene *seryl-tRNA synthetase* (PF3D7_0717700) was calculated by using the 2^−ΔΔCt^ method [125].

### RNA fluorescent in-situ hybridization (FISH)

RNA FISH was performed as described [31]. Briefly, purified ring-stage parasites were lysed with saponin and released parasites fixed in suspension with ice-cold 4% paraformaldehyde. Parasites were then deposited on Teflon coated microscope slides and hybridized with denatured *var* probes at 42℃ for at least 16 h. All the FISH probes were PCR amplified from genomic DNA using the primers listed in Table S9B. The slides were then washed three times in 2×SSC at 42℃. Finally, the slides were incubated with streptavidin-488 antibody at room temperature for 30 min. Images were taken using a Nikon ECLIPSE E600 epifluorescence microscope. NIS Elements 3.0 software was used for acquisition and ImageJ for composition.

### Statistical analysis

For all experiments, three or more independent biological replicates were performed. The results are presented as mean ± SD. Results are regarded significant if *P* < 0.05 as established by ANOVA, Fisher’s exact test, paired Mann Whitney *U* test or paired Wilcoxon test, and the respective analysis was shown in the figure legends. To analyze the schizont numbers containing different numbers of merozoites, a χ^2^ goodness of fit test was first used to evaluate if the number of schizonts that contain a certain number of merozoites was independent of the parasite lines. Then the proportions of schizonts with a certain number of merozoites were compared among these cell lines based on ANOVA for each merozoite number.

## Acknowledgments

We thank Dale Chaput for proteomic studies; Xiaolian Li for parasite culture and generation of transgenic lines. We are grateful to Jacobus Pharmaceuticals for providing the drug WR99210. This study was supported by grants U19AI 089672 and R01AI128940 from the National Institute of Allergy and Infectious Diseases, National Institutes of Health.

## Author Contributions

JM and LC designed the project. JM, LC and KK supervised the project. JM executed most experiments and interpreted data. XL performed RNA-seq experiment. SA generated the RNA-seq data and CW performed analysis of the sequence data. RJ supervised bioinformatic analysis. AL conducted IFA of H3K9Ac and chromatin immunoprecipitation, and HM performed Western blot for GCN5 expression and FISH. JM and LC wrote the manuscript. All authors read and agreed to the content of this manuscript.

## Supplementary Figures

**Figure S1.**
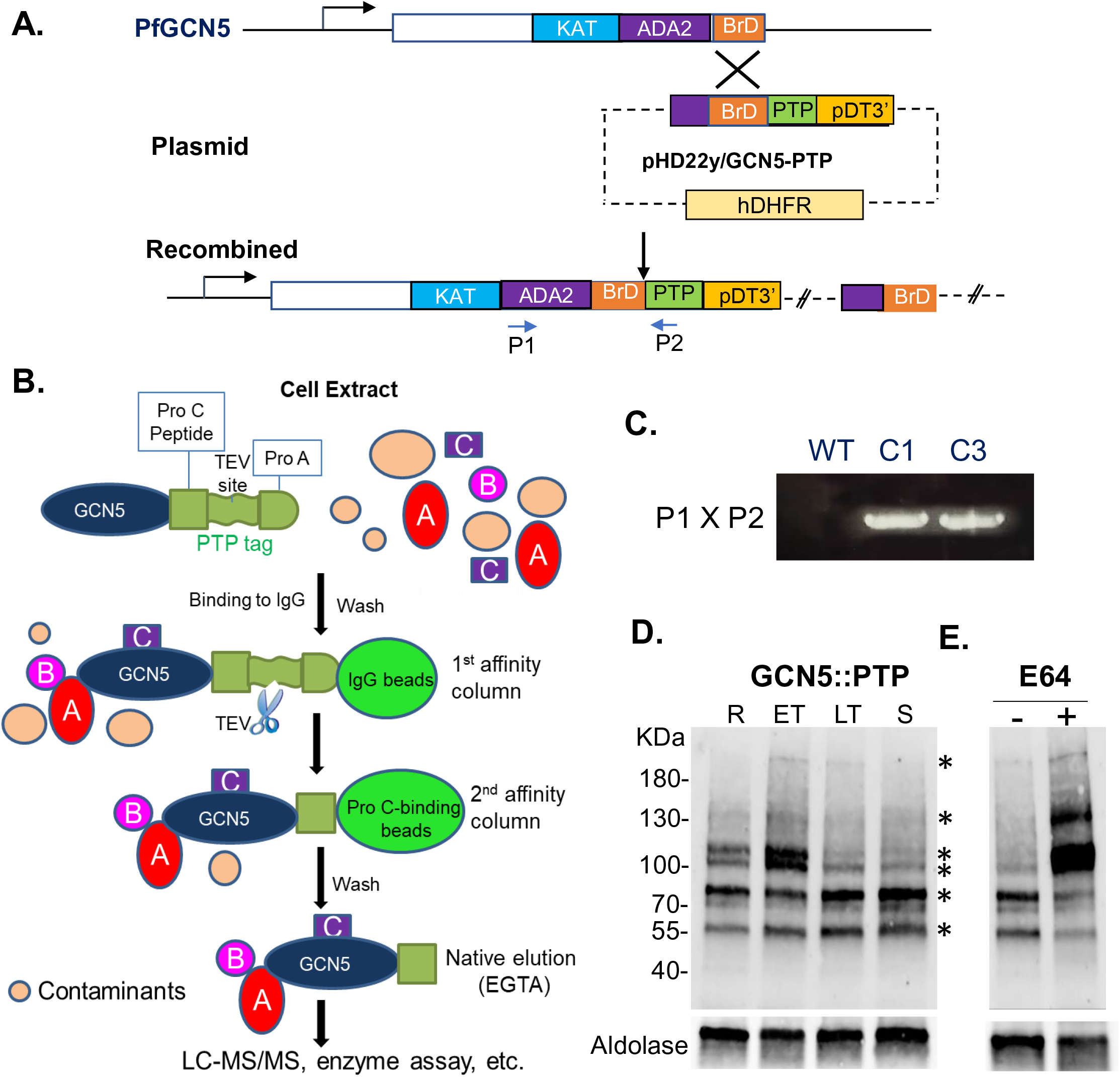
PfGCN5:PTP tagging and TAP purification. A. Schematic diagram of PTP tagging at C-terminal of PfGCN5. P1 and P2 are primers used for verification of integration by PCR. **B.** Cartoon shows the TAP procedure for purification of PfGCN5 complex. A, B, and C are subunits of the GCN5 complex. TEV: tobacco etch virus protease. **C**. PCR verification of positive clones from two clones (C1 and C3) from transfected parasite. **D**. Western blot detecting PfGCN5:PTP in the recombinant parasite clone C3 at different developmental stages (R: ring; ET: early trophozoite; LT: late trophozoite; S: schizont). The blot was probed with antibodies against protein C. Molecular markers in kDa are shown on the left. The expression of aldolase was used for equal loading control. The protein bands are indicated by asterisks. **E.** Western blot detecting PfGCN5::PTP at late trophozoite stage with or without 10 μM E64 treatment for 12 h. E64 blocks processing of PfGCN5.

**Figure S2.**
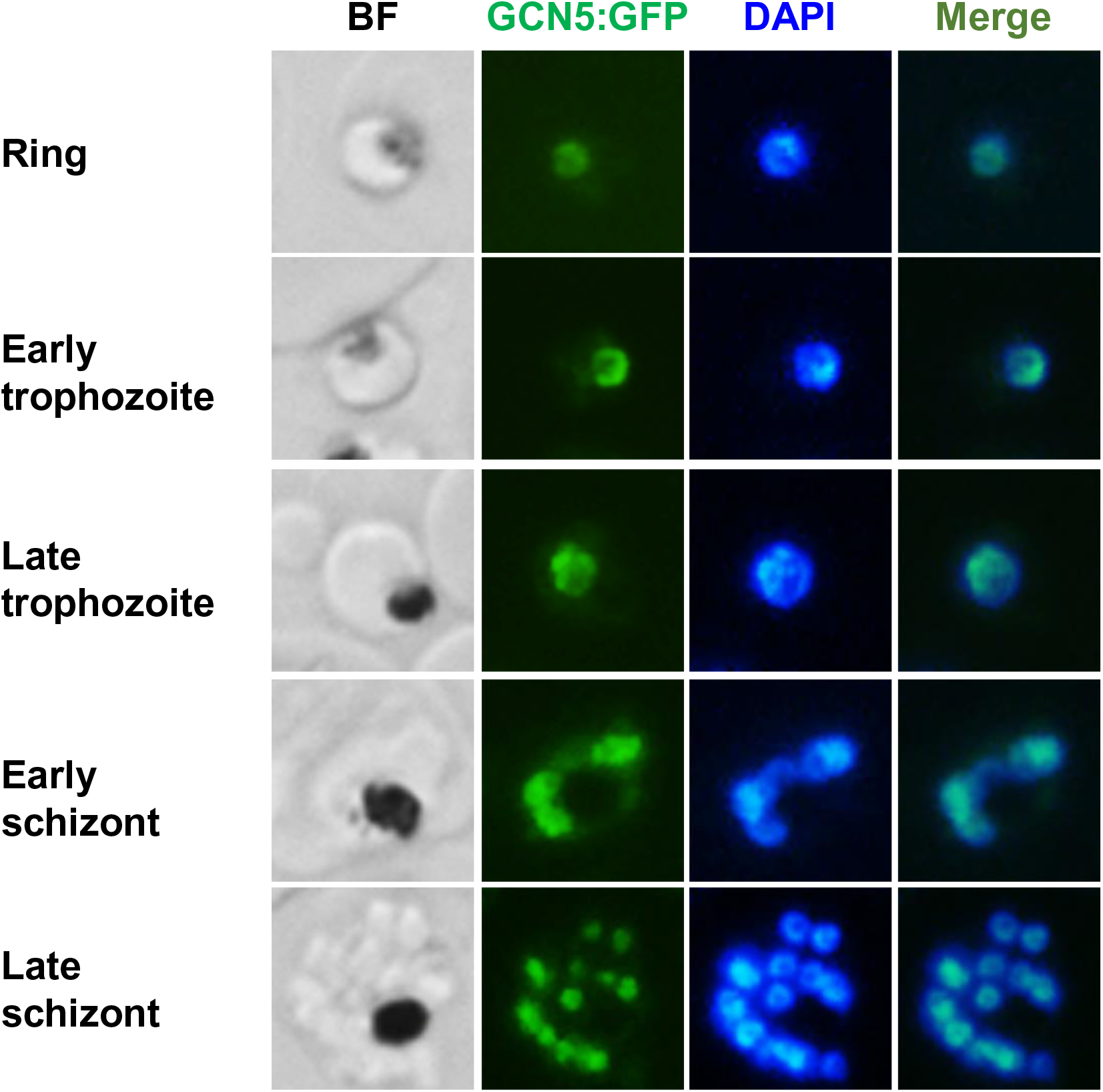
PfGCN5 expression and localization during the IDC. Live cell imaging shows the localization of PfGCN5 in GCN5::GFP parasite under a fluorescence microscope. DAPI was used to stain nucleus. BF, bright field.

**Figure S3.**
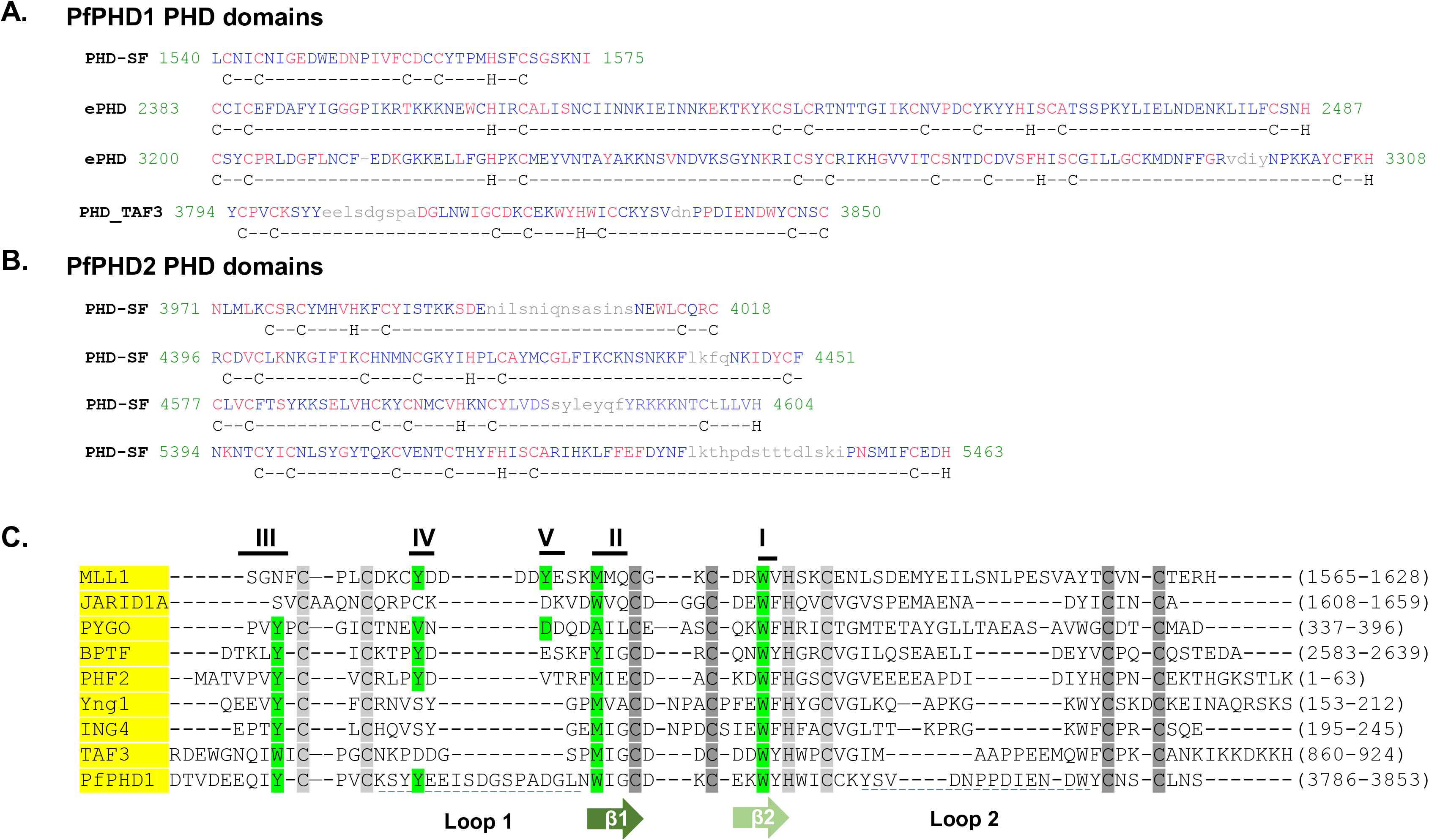
**PHD domains in PfPHD1 and PfPHD2 proteins. A**. Sequence of four PHD domains in PfPHD1, C and H amino acid residues in the PHD domain are highlighted underneath the sequence. PHD-SF: PHD superfamily; ePHD: elongated PHD domain, PHD_TAF3: TAF3 type PHD domain. **B**. Sequences of four PHD domains in PfPHD2. **C**. Alignment of PfPHD1 PHD_TAF3 domain with other known authentic PHD domains which bind H3K4me3/2. Alignment shows the conserved Zinc-binding residues in light gray for Zinc 1 and dark gray for Zinc 2, and the two core β-strands in green. The residues involved in H3K4me3 recognition are labeled I through V (forming the aromatic cages) and the aromatic residues in the recognition cage are shadowed in green. MLL1: mixed-lineage leukemia-1; JARID1A: jumonji, AT-rich interactive domain 1A; PYGO: pygopus homolog 1; BPTF: bromodomain PHD finger transcription factor; PHF2: PHD finger protein 2; Yng1: yeast homolog of mammalian ING1; ING4: inhibitor of growth protein 4; TAF3: transcription initiation factor TFIID subunit 3.

**Figure S4.**
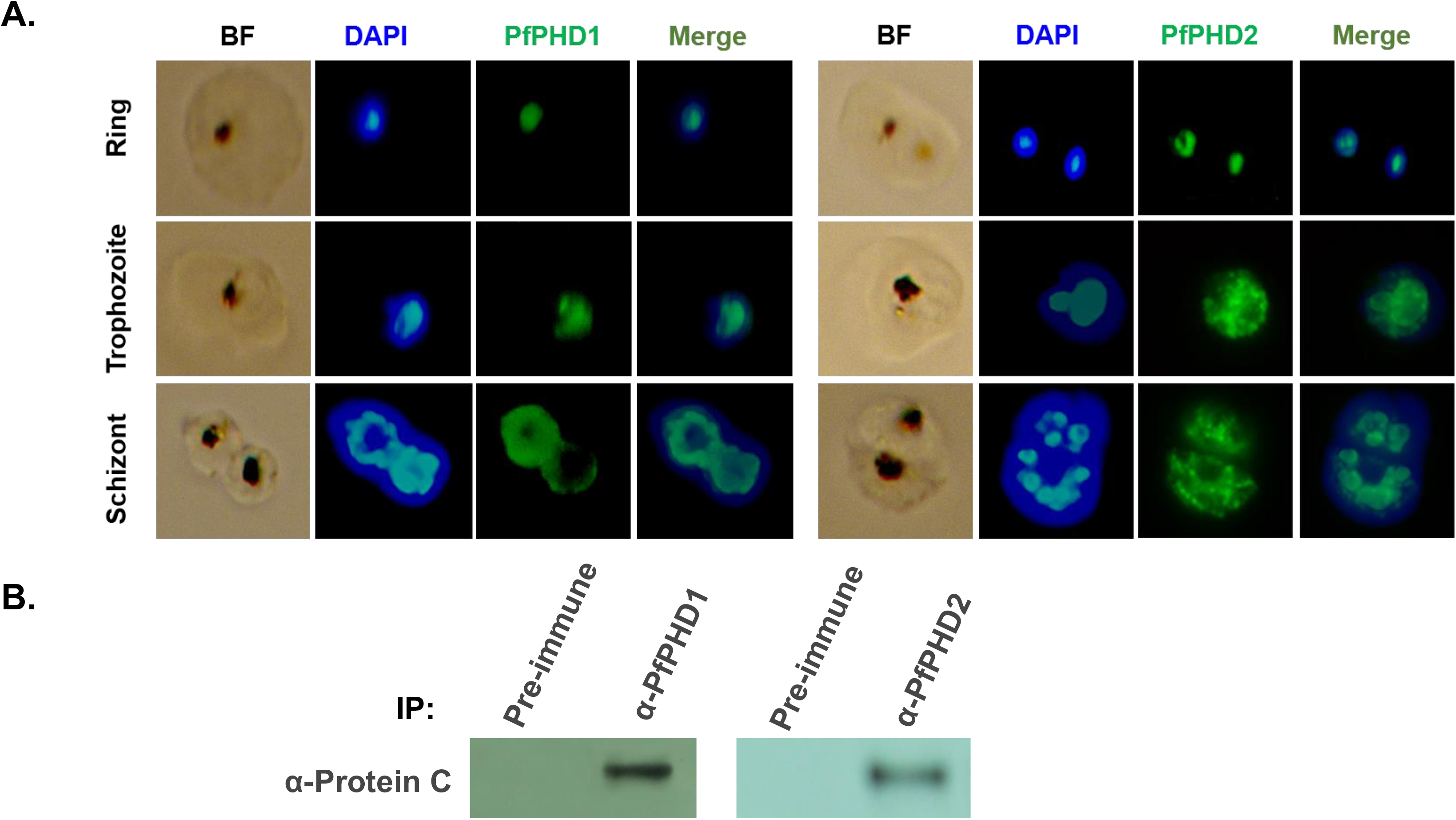
**Association of PfPHD1 or PfPHD2 with PfGCN5. A**. Nuclear localizations of PfPHD1 and PfPHD2 were interrogated by IFA using anti-PfPHD1 and PfPHD2 antibodies. Nuclei were counter-stained by DAPI. **B**. IPs of proteins from lysates of synchronized trophozoites of the PfGCN5::PTP parasite line using agarose conjugated with either anti-PfPHD1 or PfPHD2 antibodies. Immunoprecipitated proteins were separated by SDS-PAGE and probed with anti-Protein C antibodies recognizing the PTP tagged GCN5. Pre-immune sera were used as controls.

**Figure S5.**
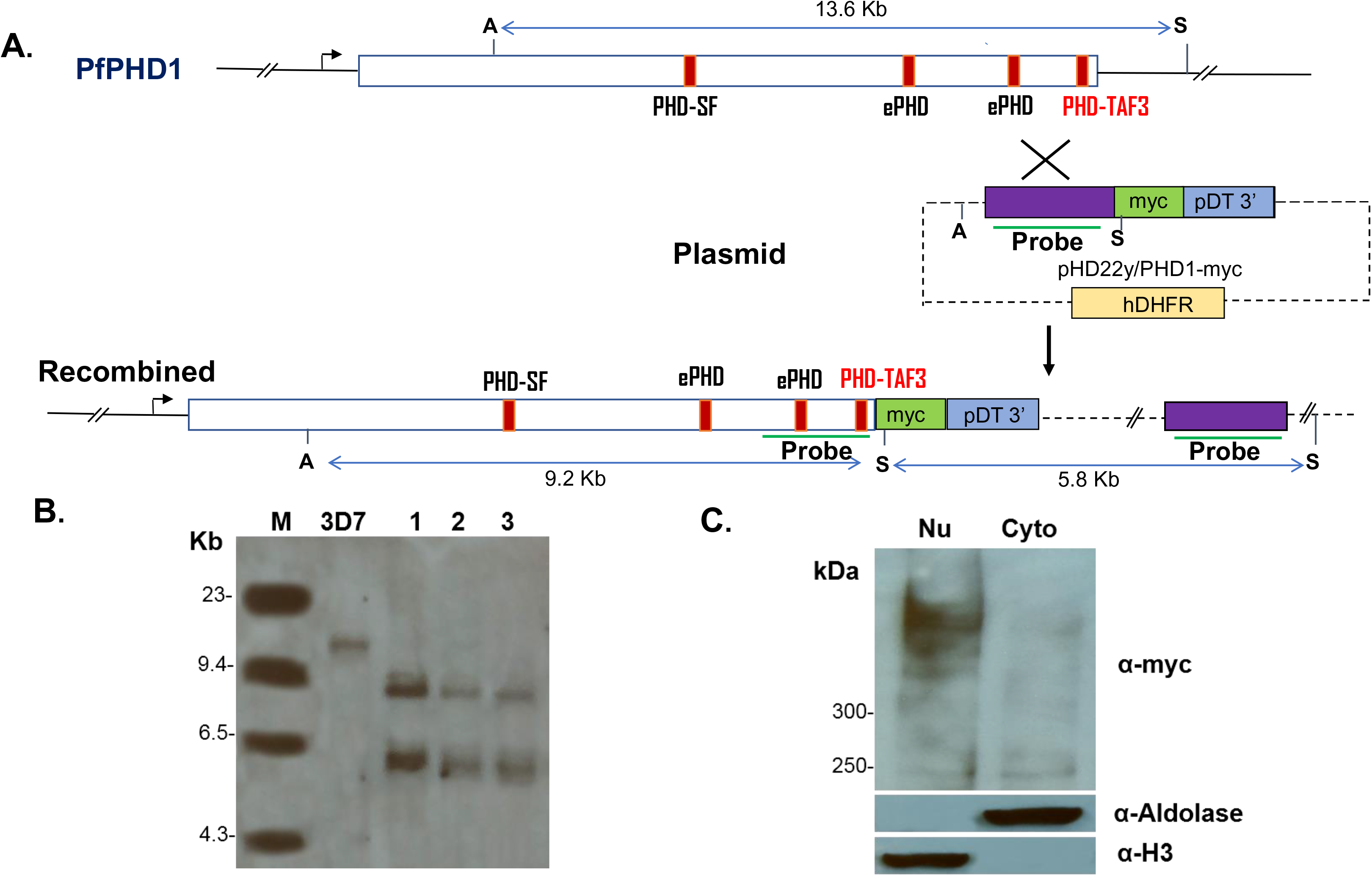
Tagging of PfPHD1 (PF3D7_1008100) with c-Myc. A. Schematic diagram of Myc tagging at C-terminal of PfPHD1. A, AvrII; S, StuI. **B.** Southern blot of 3D7 and three transgenic clones (1-3). Genomic DNA was digested with AvrII and StuI and hybridized with labeled DNA shown as “Probe” in **A**. **C.** Western blot analysis of nuclear (Nu) and cytoplasmic (Cyto) protein extracts with antibodies against the Myc tag, aldolase (for cytoplasmic compartment) and histone H3 (for nuclear compartment).

**Figure S6.**
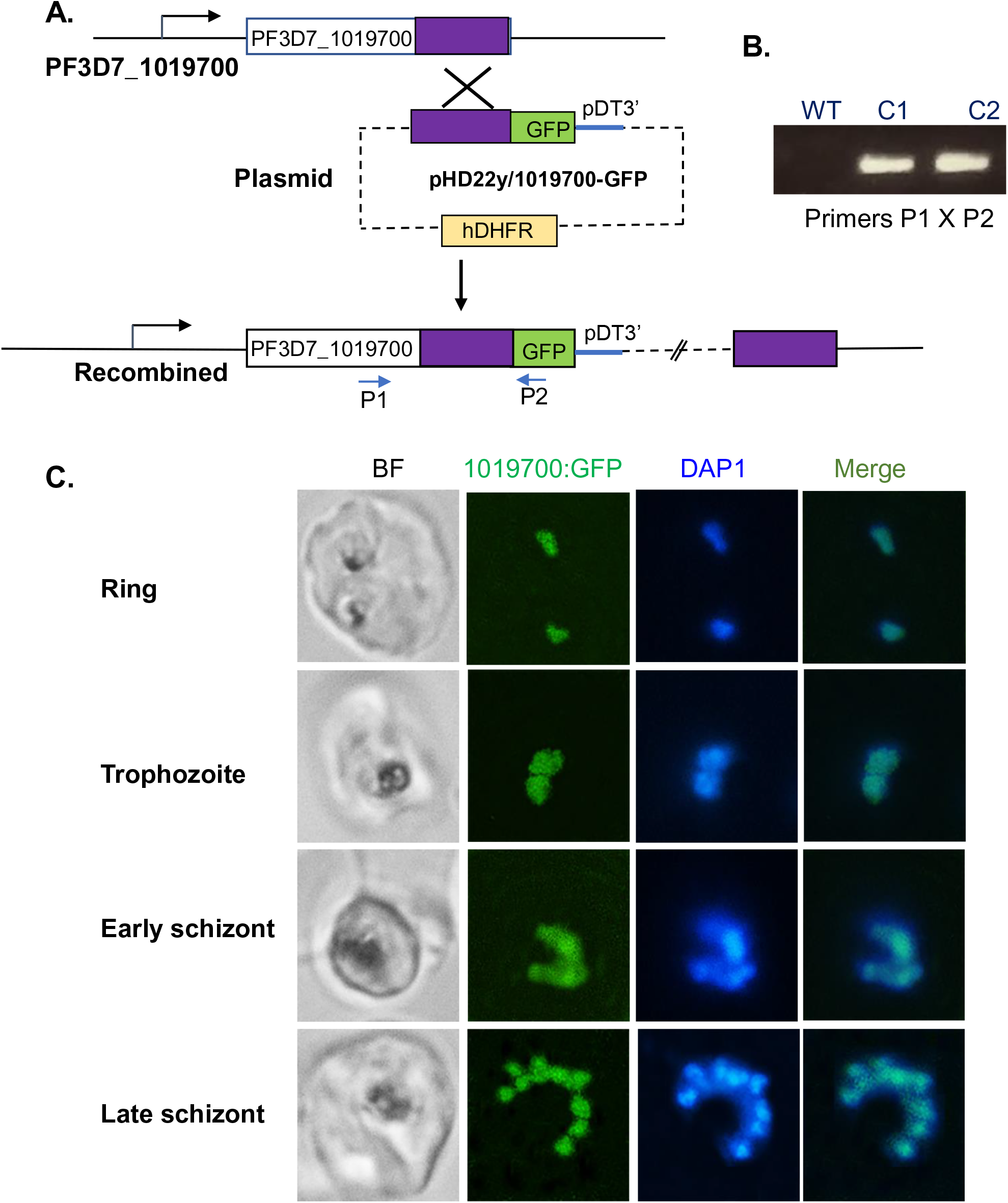
Tagging of PF3D7_1019700) with GFP. A. Diagram shows GFP tagging of the PF3D7_1019700 at its C-terminus by single-crossover homologous recombination. Purple blocks show the fragment used for homologous recombination. **B.** Integration-specific PCR using primers P1 and P2. WT, Wildtype 3D7; C1 and C2 are two transgenic clones**. C.** Live cell imaging shows the localization of PF3D7_1019700::GFP in the nuclei by fluorescence microcopy. DAPI was used to stain the nuclei.

**Figure S7.**
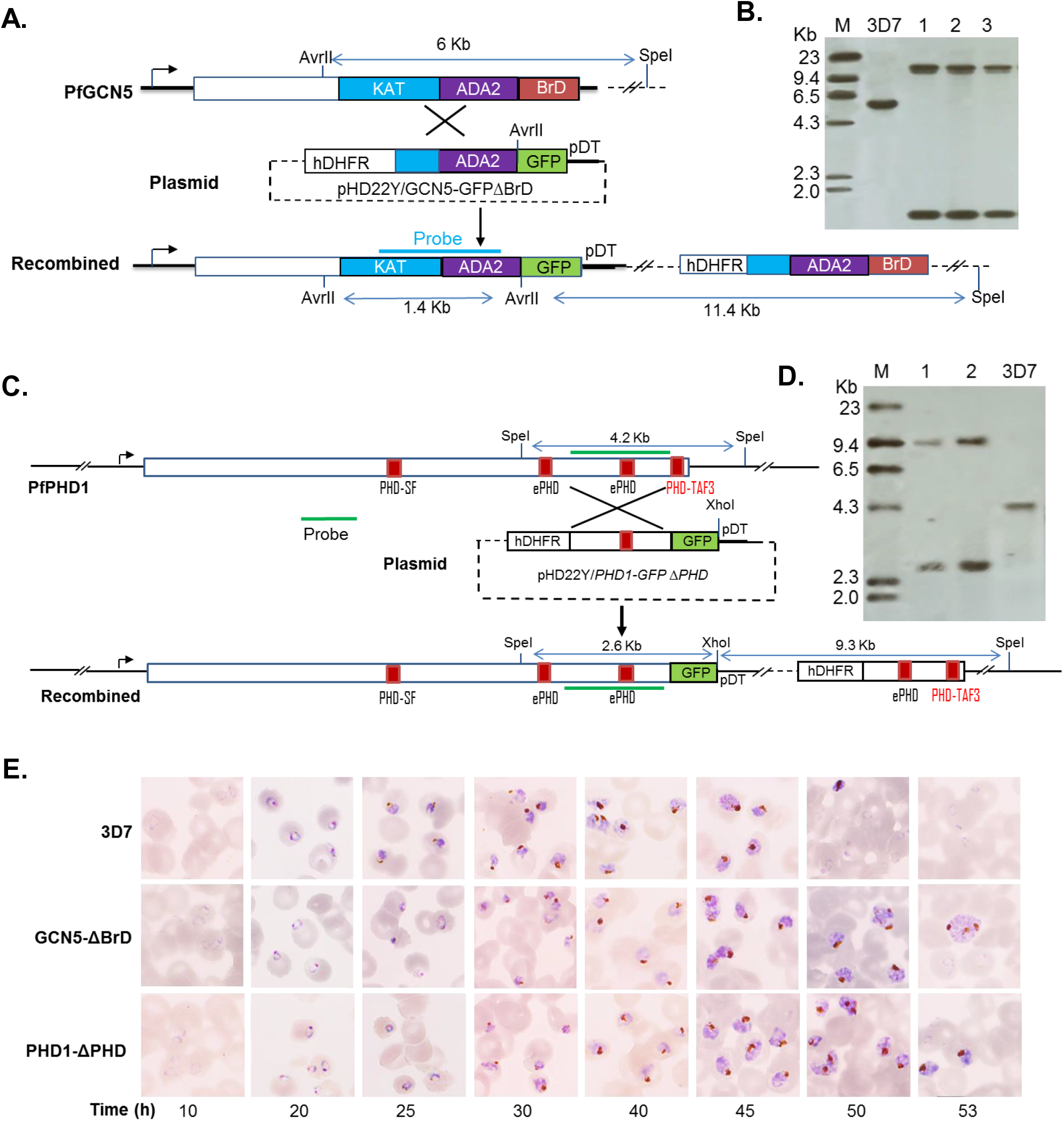
Deletion of PfGCN5 Bromodomain (BrD) and the PfPHD1 PHD-TAF3 domain. A. Schematic showing BrD deletion by single crossover homologous recombination. **B.** Southern blot analysis of three positive clones from transfected parasites. Genomic DNA was digested with AvrII and SpeI, and hybridized to the probe marked in **A**. **C.** Schematic showing the deletion of PHD-TAF3 domain. **D.** Southern blot of two positive clones from transfected parasites. Genomic DNA was digested with SpeI and XhoI, and hybridized to the probe marked in C. In both cases, GFP was tagged at the ends of truncated genes. Probes for Southern blots are marked. **E**. Images of Giemsa-stained films of parasite cultures synchronized at the ring stage to show the extended IDC of the two domain deletion parasite lines.

**Figure S8.**
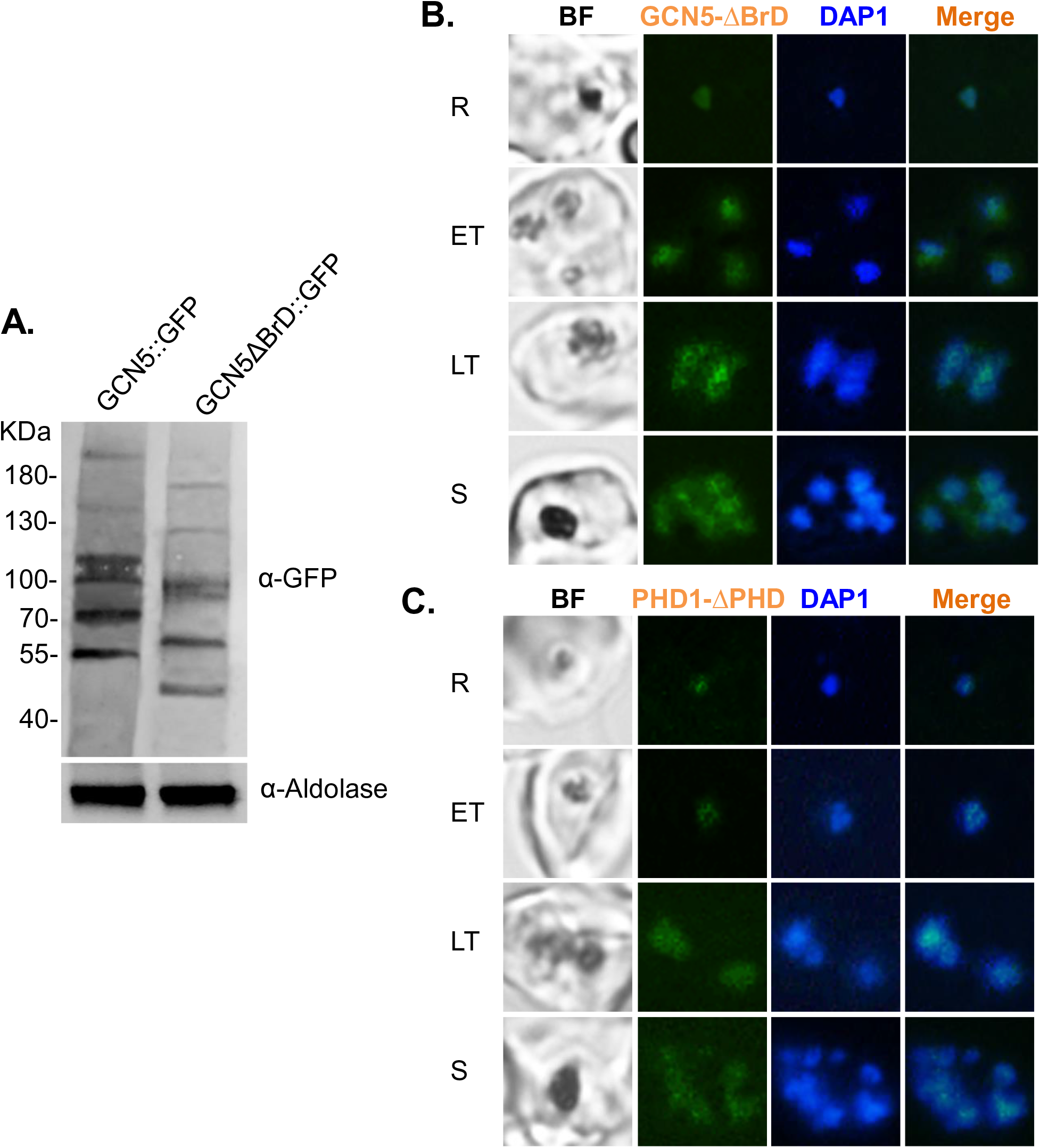
Expression of truncated PfGCN5 and PfPHD1 after domain deletions. **A**. Western blot shows the size changes of the truncated GCN5 protein bands and the reduced expression levels after BrD deletion. **B.** Live cell imaging shows GFP signals in parasite with truncated GCN5 in GCN5-ΔBrd::GFP parasite line. Compared to Figure S2, the GCN5-ΔBrd-GFP protein shows weaker fluorescence and a more diffused nuclear localization pattern. **C.** Localization of truncated PfPHD1 in PHD1-ΔPHD::GFP parasite line. Compared to Figure S4A, the PfPHD1-ΔPHD::GFP protein also shows a more diffused nuclear localization pattern.

**Figure S9.**
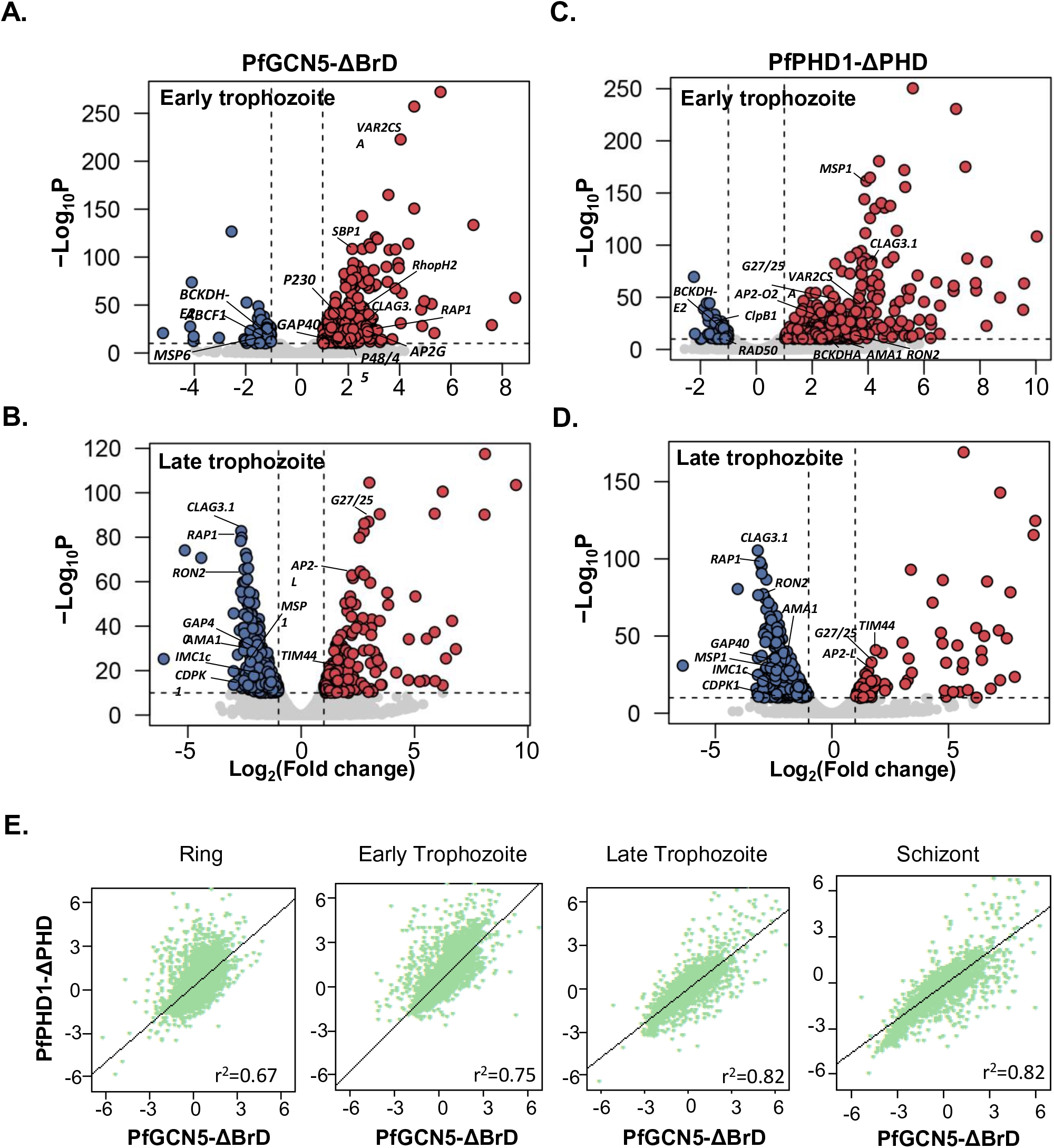
The effect of domain deletions in PfGCN5 and PfPHD1 on transcription. A–D. Volcano plots show the genes with altered transcription at the early trophozoite **(A)** and late trophozoite **(B)** stages in PfGCN5-ΔBrD, and at the early trophozoite **(C)** and late trophozoite **(D)** stages in PfPHD1-ΔPHD. **E.** Pearson correlation in fold change between PfGCN5-ΔBrD and PfPHD1-ΔPHD in different developmental stages.

**Figure S10.**
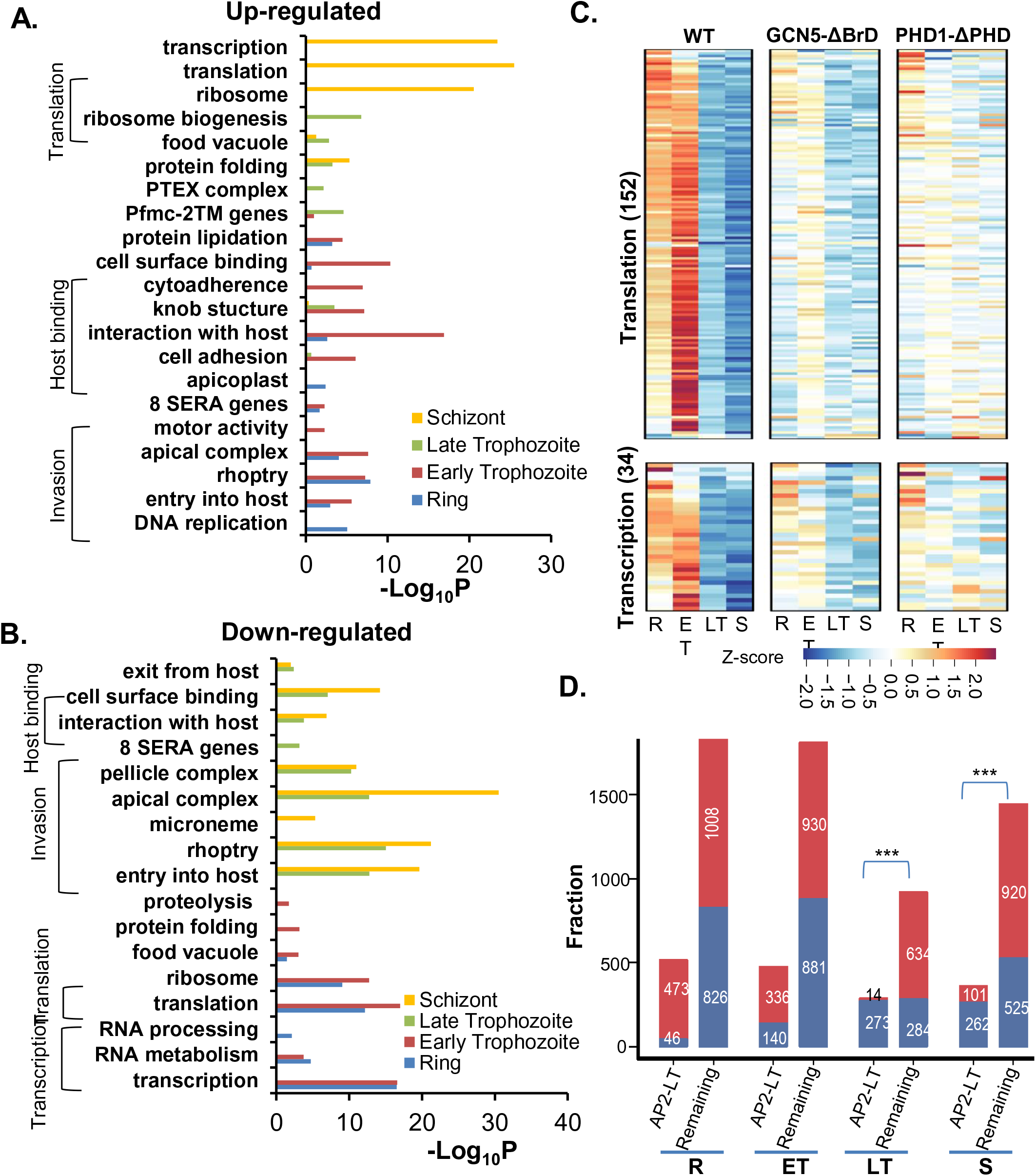
Transcriptional alteration upon domain deletions. Gene ontology enrichment analysis of up- (**A**) and down-regulated (**B**) genes in PfPHD1-ΔPHD parasites. **C.** Heatmaps display the alterations of gene transcription associated with protein translation and gene transcription upon domain deletions. **D.** Putative target genes of AP2-LT were significantly enriched in those that down-regulated in late stages of PHD1-ΔPHD::GFP. R, ring; ET, early trophozoite; LT, late trophozoite; S, schizont. ***, *P* <0.001 (Fisher’s exact test).

**Figure S11.**
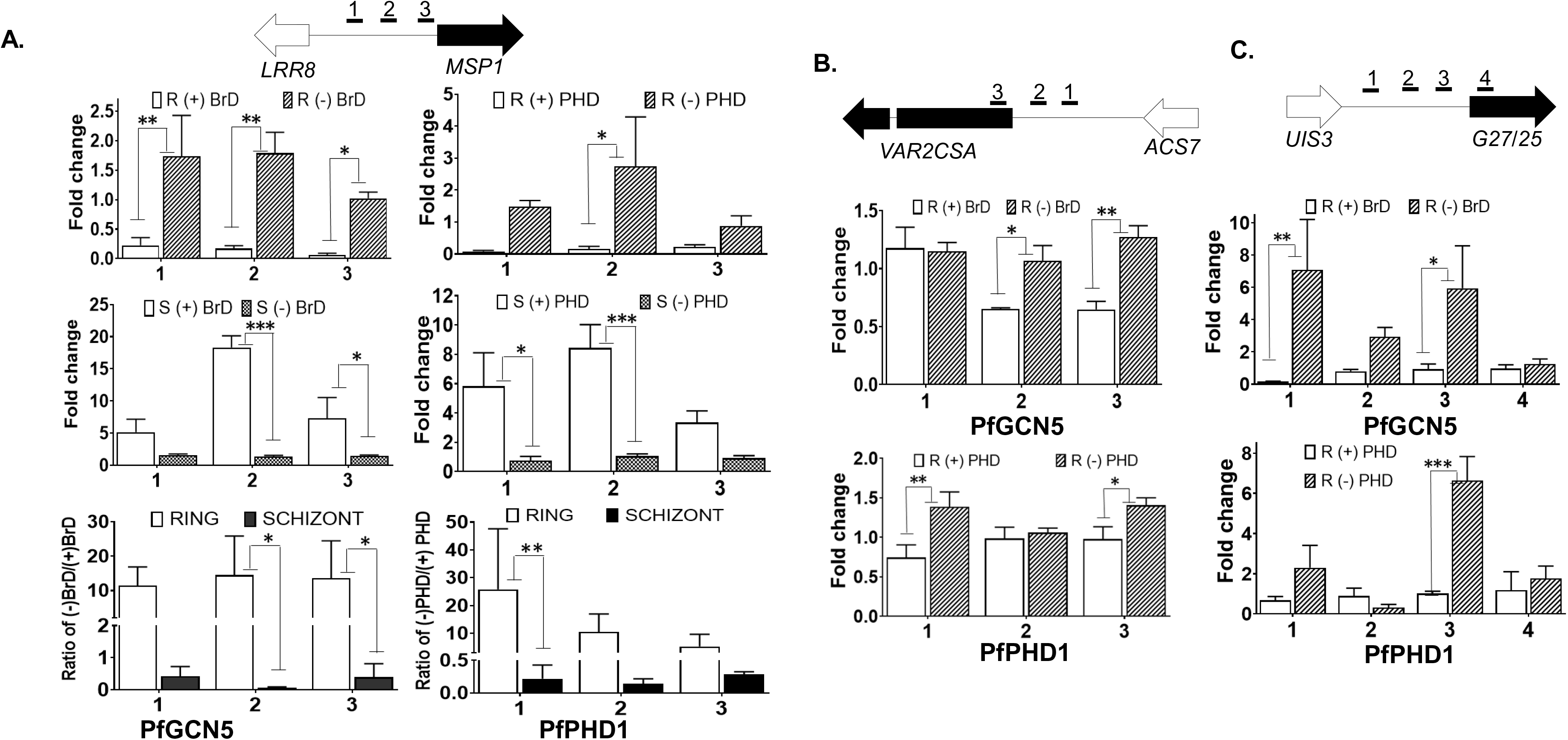
Enrichment of PfGCN5 or PfPHD1 at the promoters of genes was correlated with the activation status of the genes. The enrichment of PfGCN5 or PfPHD1 with (+) or without (−) domain deletion at the promoters of genes at the ring (R) and schizont (S) stages. **A.** *MSP1*; **B.** *VAR2CSA*; **C.** *Pfg27/25*. Enrichment was determined by chromatin-immunoprecipitation (ChIP) followed by qPCR using primer pairs marked as 1, 2, and 3 located in the promoters of the respective genes. Upon domain deletion, PfGCN5-ΔBrD and PfPHD1-ΔPHD were depleted in the promoters of *MSP1* at schizont but highly enriched at ring stage as comparing to control parasites (**A**). They were enriched at the promoters of *var2csa* (**B**) and the sexual stage gene Pfg27/25 (**C**) at the ring stage. (*, **, and *** donate P<0.05, 0.01 and 0.001, paired Mann Whitney u test)

## Supplementary Tables

**Table S1. Proteomic analyses of GCN5 associated complex**. **A**. Proteomic data from GCN5-PTP TAPs. **B**. SAINT analysis of GCN5-PTP TAPs. **C**. Proteomic data from PHD1-Myc IPs. **D**. SAINT analysis of PHD1-Myc IPs. **E**. Proteomic data from PF3D7_1019700-GFP IPs. **F**. SAINT analysis of PF3D7_1019700-GFP IPs.

**Table S2. Proteomic analyses of GCN5 associated complex after domain deletions. A**. Proteomic data from GCN5-GFP IPs. **B**. SAINT analysis of GCN5-GFP IPs. **C**. Proteomic data from GCN5-ΔBrD-GFP IPs. **D**. SAINT analysis of GCN5-ΔBrD-GFP IPs IPs. **E**. Proteomic data from PHD1-ΔPHD-GFP IPs. **F**. SAINT analysis of PHD1-ΔPHD-GFP IPs.

**Table S3. Transcriptome data of GCN5-ΔBrD::GFP line as compared to 3D7 wildtype.** Deseq2 analysis of three replicates of RNAseq data at ring (**A**), early trophozoite (**B**), late trophozoite (**C**) and schizont (**D**) stages.

**Table S4. Transcriptome data of PHD1-ΔPHD::GFP line as compared to 3D7 wildtype.** Deseq2 analysis of three replicates of RNAseq data at ring (**A**), early trophozoite (**B**), late trophozoite (**C**) and schizont (**D**) stages.

**Table S5. GO enrichment analyses of altered genes upon domain deletions. A**. GO enrichment analyses of altered genes upon BrD domain deletion in GCN5. **B**. GO enrichment analyses of altered genes upon PHD domain deletion in PHD1.

**Table S6. Transcriptional alteration of different biological pathways upon domain deletions. A**. Up-regulation of *var* gene expression at early asexual stage upon domain deletions. **B.** Down-regulation of invasion related genes upon domain deletions. **C**. Down-regulation of translation related genes upon domain deletions. **D**. Downregulation of transcription related genes upon domain deletions. **E**. Alteration of AP2 gene expression upon domain deletions.

**Table S7. Transcriptional escalation of HP1 controlled genes upon domain deletions.**

**Table S8. Transcriptional escalation of gametocyte and ookinete specific genes upon domain deletions.**

**Table S9. Primers list. A**. for tagging, domain deletion, integration checking and probe. **B**. for ChIP-qPCR and FISH

